# Integrative genetic analysis identifies FLVCR1 as an essential component of choline transport in mammals

**DOI:** 10.1101/2022.10.05.509986

**Authors:** Timothy C. Kenny, Artem Khan, Yeeun Son, Lishu Yue, Søren Heissel, Eric R. Gamazon, Hanan Alwaseem, Richard Hite, Kıvanç Birsoy

## Abstract

Genome-wide association studies (GWAS) of serum metabolites have the potential to uncover genes that influence human metabolism. Here, we combined an integrative genetic analysis associating serum metabolites to membrane transporters with a coessentiality map of metabolic genes. This analysis revealed a connection between feline leukemia virus subgroup C cellular receptor 1 (FLVCR1) - a plasma membrane protein - and phosphocholine, a downstream metabolite of choline metabolism. Loss of FLVCR1 in human cells and in mice strongly impairs choline metabolism due to a block in choline import. Consistently, CRISPR-based genetic screens identified several components of the membrane phospholipid machinery as synthetic lethal with FLVCR1 loss. Finally, cells lacking FLVCR1 exhibit mitochondrial defects and upregulate the integrated stress response (ISR) through heme regulated inhibitors kinase (HRI). Altogether, these findings identify FLVCR1 as a universal mediator of choline transport in mammals and provide a platform to discover substrates for unknown metabolite transporters.

## INTRODUCTION

Cells in multicellular organisms need a constant supply of nutrients and minerals to survive and function. Nutrient homeostasis is in part mediated by membrane carrier proteins, which facilitate the translocation of small molecule metabolites across cellular membranes. These membrane carriers comprise a functionally diverse group of proteins that differ in substrate specificity, tissue expression, subcellular localization, and topology. Consistent with the critical function of metabolites in development and growth, defects in metabolite transport have been associated with a subset of congenital disorders and diseases^1^. Furthermore, even non-pathogenic variants in transporter genes may underlie individual variability in physiologic outcomes and drug metabolism. There is increasing evidence that these carriers could also be targeted for therapy. For example, inhibitors of sodium-glucose co-transporters in the intestine are approved drugs for adults with diabetes^1^. Similarly, serotonin reuptake inhibitors are commonly used for the treatment of depression^2^. Despite their clear roles in physiology and disease, many of these small molecule carriers are poorly studied owing to their hydrophobicity. While emerging genetic, structural, and metabolomic technologies have enabled characterization of a subset of transporters, approximately 30% of them still do not have known substrates or physiological functions^3^.

Here, to link biochemical pathways to uncharacterized membrane transporter genes, we used a genome-wide association study (GWAS) of plasma metabolites from a cohort of Finnish individuals, combined with a coessentiality network of metabolic genes. This analysis identified a ubiquitously expressed plasma membrane transporter, feline leukemia virus subgroup C cellular receptor 1 (*FLVCR1)*, as a genetic determinant of phosphocholine and phosphatidylcholine levels in human plasma. Biochemical characterization of cells lacking *FLVCR1* and embryonic lethal *Flvcr1—*null mice revealed striking defects in choline metabolism. Mechanistically, we demonstrate that FLVCR1 mediates choline import in mammalian cells. Furthermore, a series of CRISPR-Cas9-based genetic screens determine membrane phospholipid metabolism as synthetic-lethal with *FLVCR1* loss. Finally, in cells lacking FLVCR1, choline deprivation results in mitochondrial dysfunction and activation of the integrated stress response (ISR) as defining features of this perturbation to cellular homeostasis. Altogether, our work identifying FLVCR1 as a key component for choline metabolism in mammals highlights the utility of uncovering novel metabolic substrates of transporters through integrative genetic analysis.

## RESULTS

### An integrative genetic analysis associates serum metabolites to membrane transporters in humans

One approach to link metabolic pathways to poorly studied small molecule transporters is to leverage serum metabolomics GWAS data and determine transporters that can impact levels of particular serum metabolites, which reflects the net production by all tissues. We therefore applied the minimum-p-value gene-based approach to the METSIM GWAS dataset of 6136 men from late-settlement Finland for which 1391 plasma metabolites were measured^4^ (Fig. 1a). Focusing on a list of 2892 metabolic genes including those that encode for enzymes and small molecule transporters, our analysis identified potential causal genes for serum levels of individual metabolites (Fig. 1a,b, S1a). Of note, associations involving xenobiotics and uncharacterized compounds were excluded from further analysis (Fig. S1b). Consistent with the critical role of plasma membrane transporters (PMTs) in nutrient homeostasis, we further restricted our analysis to 306 PMTs, of which 207 were found to have significant metabolite associations (Fig. 1b).

**Figure 1.**
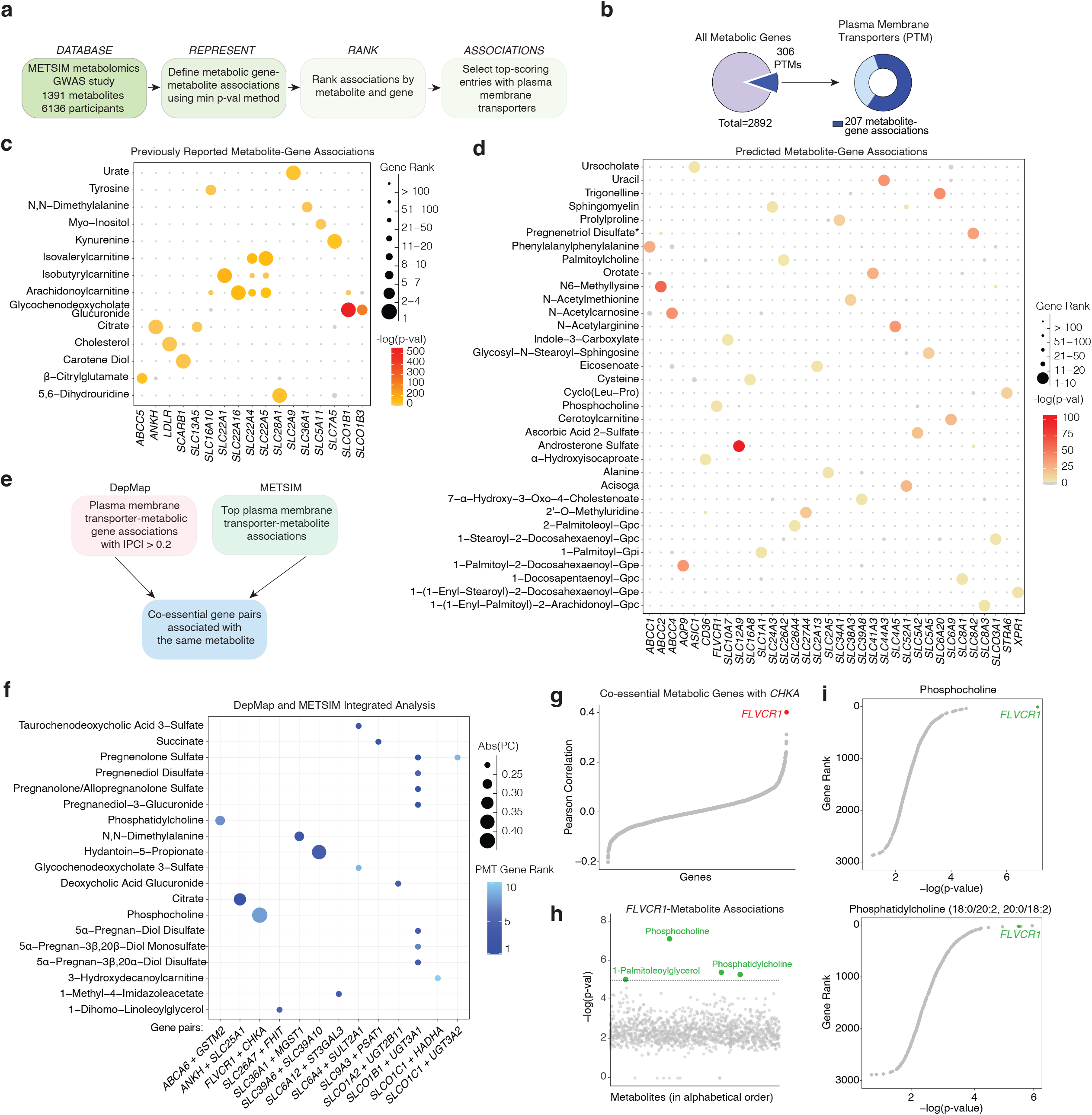
An integrative genetic analysis associates serum metabolites to membrane transporters in humans. **a)** Schematic showing the pipeline for the METSIM analysis. **b)** Summary of the METSIM analysis of 2892 metabolic genes queried for metabolite associations. Pie chart representing the number of plasma membrane transporters (PMT) for which a metabolite association was found **c)** Bubble plot of top-scoring metabolite-PMT associations previously reported. **d)** Bubble plot of undescribed metabolite-PMT associations for which the PMT had only a single significant metabolite association. Bubble color corresponds to the -log(p-val) of the association between the indicated metabolite and PMT. Bubble size represents the rank of the gene (ordered by -log(p-val) across all the analyzed metabolic genes with 1 corresponding the strongest association for the corresponding metabolite. **e)** Schematic showing the workflow of the integrated METSIM and DEPMAP analysis. **f)** Bubble plot of the integrated analysis of DepMAP and METSIM datasets. Data shown are the co-essential gene pairs with the absolute value of Pearson Correlation Coefficient (|PC|) > 0.2 computed from DepMAP CRISPR Chronos Scores that share a significant metabolite based on METSIM analysis. Bubble size represents |PC|. Bubble color corresponds to the rank of PTM for the corresponding metabolite. **g)** Pearson Correlation between *CHKA* and metabolic genes computed from CRISPR DepMAP Chronos. **h)** METSIM metabolite-*FLVCR1* associations displayed as log(p-val) of the top-scoring marker for the given metabolite within the defined *FLVCR1* genomic region vs. metabolites in alphabetical order. The dotted line indicates the significance threshold of -log(p-val) = 5. **i)** Associations between metabolic genes and phosphocholine or phosphatidylcholine (18:0/20:2, 20:0/18:2). Data are plotted as metabolic gene rank for the given metabolite vs. -log(p-val) of the top-scoring marker of the metabolite within the defined gene region.

To confirm the robustness of our approach, we first queried our dataset for previously reported metabolite-PMT associations, which yielded 17 metabolite-PMT pairs (Fig 1c). Among these were cholesterol and low-density lipoprotein receptor (LDLR), which is an established determinant of serum cholesterol levels^5^. Similarly, we identified progressive ankylosis protein (*ANKH)*, a citrate transporter and solute carrier family 7 member 5 (SLC7A5), a kynurenine importer to affect serum levels of citrate^6^, and kynurenine^7^, respectively. Our approach also revealed 132 putative associations for which a given metabolite significantly scored for one or more PMTs (Fig. 1d, Fig. S1c, Table 1). We next integrated our results with the coessentiality dataset from DepMap, which is a collection of genome-wide loss of function screens across hundreds of genomically characterized cell lines^8^. This enabled us to identify co-essential metabolic gene pairs, both of which associate with the same serum metabolite in the METSIM GWAS dataset (Fig. 1e,f). The metabolic gene pair with the strongest coessentiality was *FLVCR1* and choline kinase alpha (*CHKA)* (Pearson correlation = 0.4) (Fig. 1g). *FLVCR1* is a plasma membrane transporter previously characterized as a heme exporter^9^ and has been implicated in the rare autosomal-recessive disorder posterior column ataxia and retinitis pigmentosa (PCARP)^10–13^. *CHKA* catalyzes the production of phosphocholine through phosphorylation of choline. Importantly, in the METSIM dataset, *FLVCR1* and *CHKA* are associated with phosphocholine and its downstream metabolite phosphatidylcholine (Fig. 1h,i). Given the strong link between *FLVCR1* and choline metabolism, we focused our attention on *FLVCR1*.

### *FLVCR1* loss impairs choline metabolism in human cells and in mice

To begin to biochemically characterize the function of *FLVCR1*, we generated pairs of *FLVCR1* knockout cell lines and those complemented with *FLVCR1* cDNA, and profiled metabolites by liquid chromatography-mass spectrometry (LC-MS) (Fig. 2a, S2a,b). Among all detected metabolites, we found largest reductions in the levels of choline and its downstream metabolites phosphocholine and betaine in *FLVCR1*-knockout cells (Fig. 2a, S2b). Once imported into the cell, choline has multiple metabolic fates. The biosynthesis of phosphatidylcholine through the Kennedy pathway^14^ begins with the phosphorylation of choline by CHKA followed by a series of enzymatic reactions to generate the phosphatidylcholine, the predominant phospholipid species of cellular membranes. Additionally, choline is a key constituent of acetylcholine, a neurotransmitter molecule, and betaine, a key metabolite in one carbon metabolism. To directly determine whether production of downstream metabolites change in response to FLVCR1 loss, we performed metabolite isotope tracing experiments using choline ([1,2-^13^C_2_]Choline) in *FLVCR1*-knockout cells and those expressing *FLVCR1* cDNA (Fig. 2b), Remarkably, incorporation of isotope-labelled choline into downstream metabolites phosphocholine, betaine, and glycerophosphocholine was severely blunted in *FLVCR1*-knockout cells (Fig. 2c-e, S2c-e). Consistent with the impairment of choline metabolism, loss of FLVCR1 also led to a depletion of phosphatidylcholine (PC) species but, surprisingly, a strong accumulation of triglycerides in a lipidomic analysis (Fig. 2f, S2f). This is in line with the observation that a decrease in PC synthesis results in larger lipid droplets^15–17^. Indeed, *FLVCR1*-knockout cells accumulate significantly more lipid droplets compared to controls (Fig. 2g, S2g).

**Figure 2.**
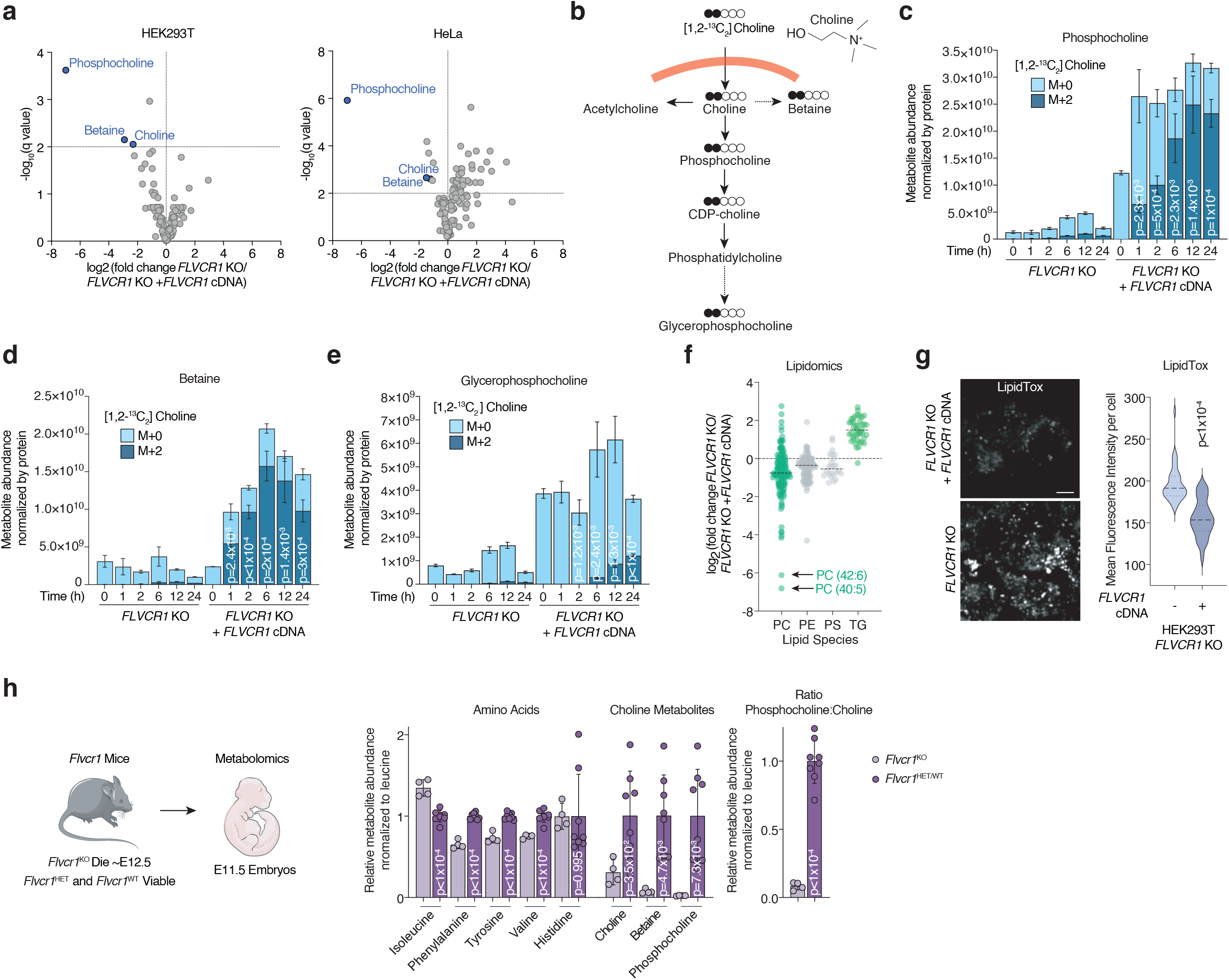
*FLVCR1* loss impairs choline metabolism in human cells and in mice. **a)** Volcano plot showing the log_2_ fold change in metabolite abundance versus -log_10_(q value) from *FLVCR1*-knockout HEK293T and HeLa cells expressing a vector control or *FLVCR1* cDNA cultured in choline depleted media for 24 hours. The dotted line indicates the significance threshold of FDR q<1% (-log_10_(q value) = 2). Statistical significance was determined by multiple two-tailed unpaired *t*-tests with discovery determined using the Two-stage linear step-up procedure of Benjamini, Krieger and Yekutieli. **b)** Molecular structure of choline and schematic for tracing of [1,2-^13^C_2_]Choline into downstream metabolites. Metabolite abundance of **c)** phosphocholine, **d)** betaine and **e)** glycerophosphocholine after incubation with [1,2-^13^C_2_]Choline for the indicated timepoints in *FLVCR1*-knockout HEK293T cells expressing a control vector or *FLVCR1* cDNA. Bars represent mean ± standard deviation; n = 3. Statistical significance determined by two-tailed unpaired *t*-test and displayed p-values compare M+2 metabolite abundance between *FLVCR1* KO and *FLVCR1* KO +*FLVCR1* cDNA at corresponding timepoints. **f)** Lipidomics analysis of *FLVCR1*-knockout HEK293T cells expressing a control vector or *FLVCR1* cDNA cultured in choline depleted media for 24 hours. Dot plot of individual lipid species of *FLVCR1*-knockout cells relative to *FLVCR1* KO +*FLVCR1* cDNA cells and grouped by lipid class – phosphatidylcholine (PC), phosphatidylethanolamine (PE), phosphatidylserine (PS), and triglyceride (TG). Dotted line represents lipid species abundance of *FLVCR1* KO +*FLVCR1* cDNA cells. Median log_2_ fold change of *FLVCR1* KO cells denoted by black line for each lipid class; n = 3. **g)** LipidTOX staining of *FLVCR1*-knockout HEK293T cells expressing a control vector or *FLVCR1* cDNA. Scale bar = 5μm. Violin plot showing quantification of mean fluorescence intensity per cell. Statistical significance was determined by two-tailed unpaired *t*-test. n = 28 cells *FLVCR1* KO; n = 39 cells *FLVCR1* KO +*FLVCR1* cDNA. **h)** Schematic of *Flvcr1* mice used for whole embryo metabolomics. Metabolite levels or ratio normalized by endogenous leucine between *Flvcr1*^KO^(n = 4) and *Flvcr1*^HET^ (n = 6) plus *Flvcr1*^WT^ (n = 2) mice. Bars represent mean ± standard deviation. Statistical significance determined by two-tailed unpaired *t*-test.

Building upon these findings in cultured cells, we next asked whether FLVCR1 loss impacts choline metabolism at the organismal level. On the C57BL/6 background, *Flvcr1*^KO^ mice are embryonic lethal with death occurring around E12.5 while *Flvcr1*^HET^ and *Flvcr1*^WT^ mice are fully viable^9^. We therefore performed whole-embryo metabolomics on E11.5 embryos from timed matings of *Flvcr1*^HET^ mice (Fig. 2h). Strikingly, we found significant drops in choline (3-fold), betaine (13-fold), and phosphocholine (43-fold) levels of *Flvcr1*^KO^ embryos when compared to *Flvcr1*^HET/WT^ controls (Fig. 2h). This is in contrast to a subset of amino acids, whose levels are unchanged or mildly different (maximum change of 50%) between *Flvcr1*^KO^ and *Flvcr1*^HET/WT^ embryos (Fig. 2h, S2h). Altogether, these results suggest that FLVCR1 is required for the maintenance of choline homeostasis in human cells and in mice.

### FLVCR1 mediates choline import and enables cell proliferation under choline limitation

We next sought to understand the precise mechanism by which FLVCR1 regulates choline metabolism in mammalian cells. Previous work identified FLVCR1 as a heme exporter and that *FLVCR1*-knockout cells accumulate heme^9,18–23^. To ascertain the relevance of these observations to our findings, we mimicked heme overload in cells with supplementation of hemin, a heme analog, followed by isotope-labelled choline tracing (Fig. S2i, S2j). Hemin did not, however, have an impact on total or labeled levels of phosphocholine or betaine (Fig. 2Sj, S2i). Given the genetic relationship between *CHKA* and *FLVCR1*, we next considered whether FLVCR1 impacts CHKA function. While FLVCR1 loss reduced all choline downstream metabolites, *CHKA*-knockout cells display a strong decrease in incorporation of isotope-labelled choline into phosphocholine, but an increase in betaine, likely due to the overflow of intracellular choline into betaine production (Fig. S2k, S2l). This raises the possibility that FLVCR1 impacts processes upstream of CHKA. Notably, *FLVCR1*-knockout cells display no observable differences in the protein abundance of enzymes involved in phosphatidylcholine synthesis (Fig. S2m).

Choline is present in human serum at ∼10μM^24^ and in extracellular fluid at ∼3-6μM^25–27^. Given that choline is an essential metabolite, most mammalian cells need to take up choline from the extracellular environment at these concentrations. A high affinity transporter, SLC5A7/CHT1, has previously been identified, which enables choline reuptake only in cholinergic neurons^28,29^(Fig. S3a). Additionally, several transporters defined for choline transport in other tissues (SLC44A1/CTL1, SLC44A2/CTL2, SLC22A1/OCT1, SLC22A2/OCT1) have much higher Km values (50-500μM) than physiological choline concentrations^30–35^. However, a high affinity choline transporter expressed across mammalian cell types has not been identified. Given the ubiquitous expression pattern of FLVCR1 and strong changes in downstream choline metabolites upon its loss (Fig. S3a), we hypothesized that FLVCR1 may mediate choline transport in most mammalian tissues.

Upon entry into the cell, CHKA phosphorylates choline to phosphocholine and traps it in the cell in a manner analogous to glucose phosphorylation by hexokinases^36^. The net accumulation of radiolabeled choline in cells is accordingly a function of both choline transport and trapping^36,37^. To test the ability of *FLVCR1* to mediate choline import, we therefore utilized radiolabeled choline ([Methyl-^3^H]Choline) uptake assays (Fig. 3a). Remarkably, loss of FLVCR1 blocked choline uptake in HEK293T and HeLa cells in a dose and time dependent manner (Fig. 3b,c Fig. S3b,c) This phenotype was completely rescued by expression of *FLVCR1* cDNA or the neuron specific high affinity choline transporter *SLC5A7*^29^ cDNA, but not by SLC44A1 cDNA in *FLVCR1*-knockout cells (Fig 3d). Similarly, phosphocholine and/or choline supplementation restored phosphocholine levels in *FLVCR1*-knockout cells (Fig. 3e, S3d). Previously reported choline transporters have been designated by their sensitivity to hemicholinium-3 inhibition and dependence on sodium for transport^31^. In contrast to *SLC5A7*, we found choline uptake via *FLVCR1* to be independent of sodium and insensitive to hemicholinium-3 (Fig S3e-i).

**Figure 3.**
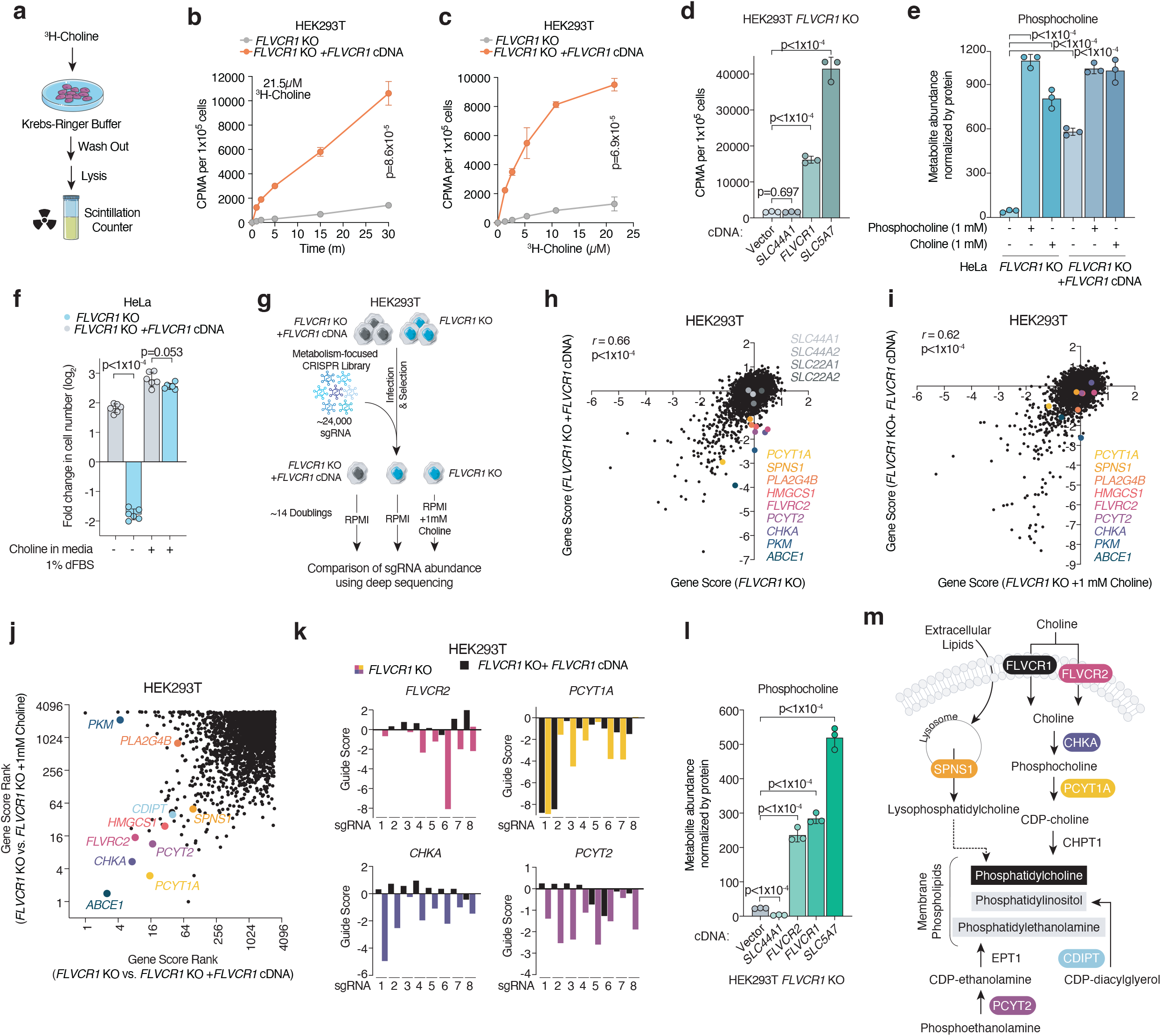
FLVCR1 mediates choline import in mammalian cells. **a)** Schematic for radioactive choline uptake assays using ^3^H-choline. **b)** Uptake of [Methyl-^3^H]-Choline in *FLVCR1*-knockout HEK293T cells expressing a vector control or *FLVCR1* cDNA incubated for indicated timepoint. Data represented as mean ± standard deviation and normalized by seeded cell number; n = 3. Statistical significance was determined by multiple two-sided unpaired *t*-tests with correction for multiple comparisons using the Holm-Sidak method. t=1 minute p=1.3×10^−3^; t =2 minutes p=2.3×10^−5^, t=5 minutes, 15 minutes, and 30 minutes p<1×10^−6^. **c)** Uptake of [Methyl-^3^H]-Choline in *FLVCR1*-knockout HEK293T cells expressing a vector control or *FLVCR1* cDNA incubated with indicated dose for 30 minutes. Data represented as mean ± standard deviation and normalized by seeded cell number; n = 3. Statistical significance was determined by multiple two-sided unpaired *t*-tests with correction for multiple comparisons using the Holm-Sidak method. 1.34μM p=6×10^−6^; 2.69μM, 5.38μM, 10.75μM, and 21.5μM p<1×10^−6^. **d)** Uptake of 21.5μM [Methyl-^3^H]-Choline in *FLVCR1*-knockout HEK293T cells expressing an empty vector (EV) control, *SLC44A1, FLVCR1*, or *SLC5A7* cDNA for 30 minutes. Data represented as mean ± standard deviation and normalized by seeded cell number; n = 3. Statistical significance determined by two-tailed unpaired *t*-test. **e)** Phosphocholine abundance in *FLVCR1*-knockout HeLa cells expressing a vector control or *FLVCR1* cDNA after incubation with 1mM phosphocholine or 1mM choline for 24 hours. Bars represent mean ± standard deviation; n = 3. Statistical significance determined by two-tailed unpaired *t*-test. **f)** Log_2_ fold change in cell number of *FLVCR1*-knockout HeLa cells expressing a vector control or *FLVCR1* cDNA grown in choline depleted or choline replete media supplemented with 1% dialyzed FBS (dFBS). Bars represent mean ± standard deviation; n = 6. Statistical significance determined by two-tailed unpaired *t*-test. **g)** Schematic of the metabolism-focused CRISPR genetic screens in *FLVCR1*-knockout HEK293T cells expressing a vector control or *FLVCR1* cDNA cultured with or without 1mM choline supplementation. **h)** CRISPR gene scores in *FLVCR1*-knockout HEK293T cells expressing a vector control (*y*-axis) or *FLVCR1* cDNA (*x*-axis). Top scoring hits color coded and previously reported potential choline transporters highlighted in gray. Pearson correlation coefficient, two-sided. **i)** CRISPR gene scores in *FLVCR1*-knockout HEK293T cells expressing a vector control and supplemented with 1mM choline (*y*-axis) or *FLVCR1* cDNA (*x*-axis). Pearson correlation coefficient, two-sided. **j)** Comparison of gene score ranks from CRISPR screens. *FLVCR1*-knockout HEK293T cells expressing a vector control vs. *FLVCR1* cDNA (*x*-axis) and *FLVCR1*-knockout HEK293T cells expressing a vector control grown with or without 1mM choline supplementation (*y*-axis). **k)** Individual sgRNA scores targeting *FLVCR2, PCYT1A, CHKA*, or *PCTY2* in *FLVCR1*-knockout HEK293T cells expressing a vector control (color) or *FLVCR1* cDNA (black). **l)** Phosphocholine abundance in *FLVCR1*-knockout HEK293T cells expressing an empty vector (EV) control, *SLC44A1, FLVCR2, FLVCR1*, or *SLC5A7* cDNA. Bars represent mean ± standard deviation; n = 3. Statistical significance determined by two-tailed unpaired *t*-test. **m)** Schematic of phospholipid synthesis reactions and salvage pathway. Scoring genes from CRISPR screens which are essential in *FLVCR1*-knockout cells and rescued with choline supplementation are highlighted with color.

We next asked whether FLVCR1 is required for mammalian cells to proliferate under choline limitation. To address this, we cultured FLVCR1-null HEK293T and HeLa cells expressing a vector control or *FLVCR1* cDNA in choline depleted or replete media over multiple passages and found a subtle, albeit significant, impairment in proliferation of *FLVCR1*-knockout cells in choline depleted media conditions (Fig. S3j,k). Given that trace choline amounts may still be detectable in dialyzed FBS (dFBS), we further performed the proliferation assays under 1% dFBS and saw a marked decrease in the growth of *FLVCR1*-knockout cells which can be restored by choline repletion (Fig. 3f, S3k). These results suggest that FLVCR1 mediates choline import, and that *FLVCR1*-mediated choline uptake is essential for the growth of mammalian cells.

### Metabolic genes essential for cell proliferation upon FLVCR1 loss

FLVCR1 loss did not impact cell proliferation under standard media conditions, raising the possibility that other metabolic enzymes or transporters may compensate for its loss. To identify genes essential for cellular proliferation in the absence of *FLVCR1*, we performed CRISPR-Cas9-based genetic screens using a metabolism-focused single guide RNA (sgRNA) library^38^ in the presence and absence of supraphysiological choline^40^(Fig. 3g). Top scoring genes from these screens were those involved in the biosynthesis of phospholipids (*CHKA, PCYT2, PCYT1A, CDIPT*), whose essentiality are completely abrogated upon choline supplementation (Fig. 3h-m, S4a-d). Additionally, a recently discovered lysosomal lysophospholipid transporter^39^, *SPNS1*, scored as differentially essential in *FLVCR1*-knockout cells and choline dependent. SPNS1 mediates the transport of lysophosphotidylcholine, a lysosomal breakdown product of extracellular lipids, into the cytosol thereby providing a source for choline upon extracellular choline limitation. Consistently, lysophosphatidylcholine lipid species accumulate in the lysosome upon *SPNS1* knockout which can be detected by whole-cell lipidomics (Fig S4d). This strongly suggests that the lysosomal phospholipid salvage pathway enables cell survival when choline is depleted from proliferating cells. Other scoring genes include *HMGSC1* – an enzyme which produces the cholesterol precursor mevalonate, *ABCE1* – a cotranslational quality control factor associated mitochondrial out membrane-localized mRNA^40^ and cytosolic phospholipase *PLA2G4B*^41,42^ (Fig. 3h-m, S4a-c).

Of note, none of the previously reported choline transporters (*SLC44A1, SLC44A2, SLC22A1*, and *SLC22A2*) scored in *FLVCR1*-knockout cells (Fig. 3h, S4a). These results are consistent with the essential role of FLVCR1 in phosphatidylcholine metabolism. Additionally, *FLVCR2*, a paralog of *FLVCR1* with high sequence homology (Fig. S4e), scored as essential in *FLVCR1*-knockout cells in a choline dependent manner (Fig. 3h-m, S4a-c). Like its paralog, *FLVCR2* is ubiquitously expressed across human tissues (Fig. S4f). These data suggest that *FLVCR2* takes up choline in a compensatory fashion upon *FLVCR1* loss. To address this, we performed metabolomics assays on *FLVCR1*-knockout cells complemented with *FLVCR2, FLVCR1, SLC5A7* and *SLC44A1* cDNAs. These experiments showed that expression of FLVCR2, but not SLC44A1, restored the decrease in phosphocholine levels upon *FLVCR1* loss (Fig. 3l). In line with these results, expression of *FLVCR2* in *FLVCR1*-knockout cells partially restored radiolabeled choline uptake (Fig. S4f).

### FLVCR1-mediated choline uptake is required for mitochondrial homeostasis

We next sought to determine the physiological consequences of cellular choline depletion upon FLVCR1 loss in mammalian cells. To address this in an unbiased manner, we performed RNA-Seq analysis on *FLVCR1*-knockout and control HeLa cells after acute exposure to choline depleted or replete media. Remarkably, only *FLVCR1*-knockout cells displayed a distinct transcriptional response to choline depletion, highlighting a synergy between *FLVCR1* loss and choline limitation (Fig. 4a,b). Gene set enrichment analysis for upregulated transcripts revealed a strong enrichment for the integrated stress response (ISR) including amino acid deprivation, unfolded protein response, endoplasmic reticulum stress, and ATF4-activated gene expression (Fig. 4c). Given that choline phosphorylation is the rate limiting enzymatic step of phosphatidylcholine synthesis^36,37^, we next asked whether a decrease in phosphocholine levels may phenocopy ISR activation by *FLVCR1* loss (Fig. S5a,b). To test this, we targeted *CHKA* genetically by CRISPR or pharmacologically using a small molecule inhibitor. Similar to FLVCR1 loss, inhibition of CHKA resulted in ISR activation as measured by ATF4 and CHOP immunoblotting (Fig. S5c,d). Of note, this effect was blunted by supplementation of phosphocholine (Fig. S5d). The ISR is mediated by eIF2α phosphorylation by four distinct kinases (HRI, PERK, PKR, PERK) which are activated by diverse perturbations ^41^. To identify the kinase responsible for ISR activation under choline limitation, we generated mixed population knockout cells of all four eIF2α kinases. Among these, loss of *HRI*, but not others, impaired induction of the ISR upon CHKA inhibition (Fig. 4d). HRI has recently been shown to be activated by mitochondrial stress via DELE1^44–47^, a protein that translocates from the mitochondria to the cytosol where it binds and activates HRI^42,43^. Interestingly, *DELE1* loss also blunted ISR activation (Fig. 4e, S5e,f), indicating that impairment of choline utilization activates ISR through the mitochondrial stress response pathway.

**Figure 4.**
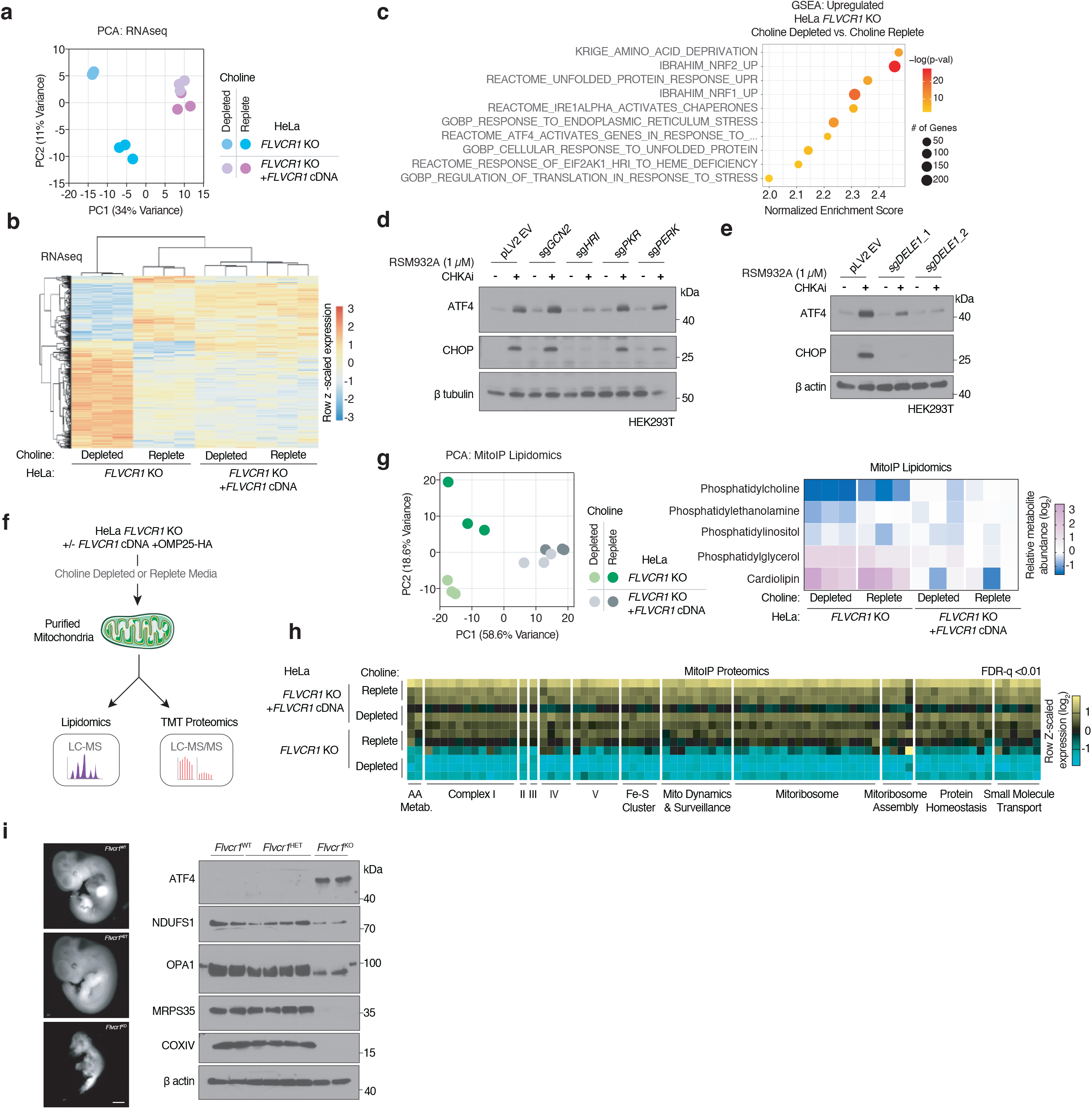
FLVCR1-mediated choline uptake is required for mitochondrial homeostasis. **a)** Principal component analysis (PCA) of RNA-Seq from *FLVCR1*-knockout HeLa cells expressing a vector control or *FLVCR1* cDNA cultured in choline depleted or choline replete media for 24 hours. **b)** Heat map of differentially expressed genes from RNA-Seq of *FLVCR1*-knockout HeLa cells expressing a vector control or *FLVCR1* cDNA cultured in choline depleted or choline replete media for 24 hours. **c)** Gene set enrichment analysis (GSEA) of significantly upregulated transcripts in *FLVCR1*-knockout HeLa cells expressing an empty vector control in choline depleted media vs. choline replete media. **d)** Immunoblotting of indicated proteins in HEK293T cells expressing an empty vector control (pLV2 EV) or sgRNA targeting *GCN2, HRI, PKR*, or *PERK* and treated with CHKA inhibitor RSM932A (1μM) or vehicle for 24 hours. **e)** Immunoblotting of indicated proteins in HEK293T cells expressing an empty vector control (pLV2 EV) or two different sgRNA targeting *DELE1* and treated with CHKA inhibitor RSM932A (1μM) or vehicle for 24 hours. **f)** Schematic for proteomic and lipidomic profiling of mitochondria from *FLVCR1*-knockout HeLa cells expressing an empty vector control or *FLVCR1* cDNA and after extended culture in choline depleted and replete conditions. **g)** Principal component analysis (PCA) of mitochondrial lipidomics from *FLVCR1*-knockout HeLa cells expressing a vector control or *FLVCR1* cDNA. Heatmap of mitochondrial lipid species. **h)** Heatmap of most differentially expressed mitochondrial proteins organized by ontology from *FLVCR1*-knockout HeLa cells expressing a vector control or *FLVCR1* cDNA cultured in choline depleted or choline replete media. Statistical significance was determined by an ANOVA test with a permutation-based FDR q<1% considered significant. **i)** Representative images of embyros of indicated genotypes at E11.5. Scale bar = 1mm. Immunoblotting of indicated proteins in *Flvcr1*^WT^, *Flvcr1*^HET^, and *Flvcr1*^KO^ whole E11.5 embryos.

To further explore how choline depletion influence mitochondria, we immunopurified mitochondria from *FLVCR1*-knockout HeLa cells expressing a vector control or *FLVCR1* cDNA using a previously described mitochondrial purification method^44^ and performed quantitative lipidomics and proteomics on these cells after exposure to choline depleted or replete conditions (Fig. 4f, S5g). Phosphatidylcholine is the most abundant phospholipid of the mitochondrial membranes. Despite this, mitochondria lack the machinery to synthesize it, and rely on the transport of PC from other cellular membranes. Consistent with this dependency, we observed a decrease in mitochondrial PC levels from *FLVCR1*-knockout cells compared to other phospholipids (Fig. 4g). Building upon our RNA-Seq results, we next hypothesized that PC depletion may have a substantial impact on mitochondrial homeostasis. Indeed, quantitative mitochondrial proteomics identified significant changes in the mitochondrial proteome of *FLVCR1*-knockout cells specifically under choline deprivation (Fig. 4h, S5h). Many fundamental mitochondrial processes and components were significantly downregulated in *FLVCR1*-knockout cells under choline deprivation including electron transport chain complexes, iron-sulfur cluster biogenesis, the mitochondrial ribosome, and protein homeostasis (Fig. 4h). These results suggest extensive mitochondrial dysfunction in cells lacking *FLVCR1* and deprived of choline. Consistently, when we compared our proteomic dataset to a list of mitochondrial proteins downregulated across diverse insults to mitochondrial homeostasis^45^, we found a significant overlap with proteins depleted in *FLVCR1*-knockout cells (Fig. S4i). Building on these observations, we next asked whether *Flvcr1*^KO^ E11.5 embryos exhibit defects similar to those observed in culture conditions. Remarkably, loss of FLVCR1 in mice strongly upregulates ATF4 protein and downregulates levels of several mitochondrial proteins. Of note, we observed ∼18 fold reduction in aspartate in levels in *Flvcr1*^KO^ embryos (Fig. 2Sh), a metabolic phenotype previously linked to mitochondrial dysfunction^46,47^. Altogether, our data implicate a functional link between choline deprivation and ISR activation through the mitochondria mediated by DELE1 and HRI.

## DISCUSSION

Choline is an essential small molecule with diverse roles in neurotransmitter synthesis, membrane homeostasis and one-carbon metabolism. Indeed, dietary choline deprivation has been associated with liver dysfunction, neurological disorders, and muscle damage^48^. Given that choline is a charged molecule, cells need to transport it from the serum and extracellular fluid via dedicated membrane carriers. In this study, we propose *FLVCR1* and its paralog *FLVCR2* as the major mediators of choline import in mammals. Our data in human cells suggest that the vast majority of intracellular phosphocholine is generated from the phosphorylation of imported choline. FLVCR1-mediated choline uptake is essential for life as its complete loss leads to embryonic lethality, suggesting that the salvage pathway cannot completely compensate *in vivo*. Similarly, patients with missense mutations in *FLVCR1* present with posterior column ataxia and retinitis pigmentosa (PCARP), an autosomal recessive neurodegenerative syndrome characterized by loss of retinal function and subsequent degeneration of the posterior columns of the spinal cord from proprioception loss^10–13^. Interestingly, the photoreceptor cells of the retina have high affinity for choline and require an abundance of phospholipids to maintain the large membranous surface area of the outer segment^49–51^. Choline availability has also been linked to retinal degeneration^52–54^. Further confirming our results, the connection between *FLVCR1* and choline was recently described as a metabolite quantitative trait loci association in nonagenarians and centenarians^55^. Our findings therefore suggest that patients with mutations in *FLVCR1* may benefit from choline supplementation, a well-tolerated dietary intervention^56^. Of note, previous work identified *FLVCR1* as a heme export protein and attributed the embryonic lethality of *Flvcr1*-null mice to ineffective erythropoeisis^9^. However, we failed to identify a transcriptional or genetic signature association with heme accumulation in our assays. Instead, our results show that *FLVCR1* depletion causes strong changes in the mitochondrial proteome and lipidome, linking choline availability to mitochondrial homeostasis. While the precise mechanism for this is unclear, phosphatidylcholine depletion may impact the stability of the TIM23 complex, the presequence translocase necessary for transport of mitochondrially targeted proteins into or across the mitochondrial inner membrane^57^. Further work is needed to decipher these connections and establish whether mitochondrial dysfunction occurs as a consequence of mitochondrial lipid remodeling under choline deprivation. Such future endeavors should also identify the molecular players involved in regulating FLVCR1 function and the structural mechanism of choline transport.

## ACKNOWLEDGEMENTS

We thank all members of the Birsoy laboratory for helpful suggestions. We thank Janis L. Abkowitz for generously providing the *Flvcr1* mice. Additionally, we thank Michael Boehnke for providing the METSIM dataset. Data was generated by the Proteomics Resource Center (RRID:SCR_017797) and the Flow Cytometry Resource Center at The Rockefeller University. Some figures use modified illustrations from Servier Medical Art licensed under a Creative Commons Attribution 3.0 Unported License.

T.K. is supported by NIH/NIDDK (F32 DK127836), the Shapiro-Silverberg Fund for the Advancement of Translational Research and a Merck Postdoctoral Fellowship at The Rockefeller University. K.B. is supported by the NIH/NIDDK (R01 DK123323-01), Mark Foundation Emerging Leader Award; and is a Searle and Pew-Stewart Scholar.

## AUTHOR CONTRIBUTIONS

Conceptualization, K.B. and T.K.; Methodology, T.K., A.K., H.A., and K.B.; Formal Analysis, T.K., A.K.; Investigation, T.K, A.K., L.Y., Y.S., S.H., E.G., H.A., R.H.; Writing – Original Draft, T.K. and K.B.; Funding Acquisition, T.K. and K.B.

## DECLARATION OF INTERESTS

K.B. is scientific advisor to Nanocare Pharmaceuticals and Atavistik Bio. Other authors declare no competing interests.

## Figure Legends

**Supplemental Figure 1. Analysis of METSIM GWAS enables prediction of unknown substrates for plasma membrane transporters. a)** Heatmap of associations between 2892 metabolic genes and 1391 metabolites identified by the minimum p-value algorithm run on METSIM GWAS data. Color gradient corresponds to gene rank for the given metabolite (ranked by p-val with 1 and 2892 corresponding to the strongest and weakest associations, respectively). Dendrogram grouping of the genes represents a hierarchical clustering on row. Metabolite class/group is color-coded and metabolites of each class/group are plotted together in columns. **b)** Bubble plot of associations between plasma membrane transporters (PMT) and xenobiotics, partially characterized, or uncharacterized compounds. **c)** Bubble plot of previously unreported PMT-metabolite associations for which a metabolites scores for more than one PMT. Bubble color corresponds to the -log(p-val) of the association between the indicated metabolite and PMT. Bubble size represents the rank of the gene (ordered by -log(p-val) across all the analyzed metabolic genes with 1 being the strongest association for the corresponding metabolite.

**Supplemental Figure 2. FLVCR1 dependent choline metabolism cannot be explained by heme overload or expression of phospholipid synthesis enzymes. a)** Immunoblotting of indicated proteins in *FLVCR1*-knockout HEK293T and HeLa cells expressing a vector control or *FLVCR1* cDNA. **b)** Volcano plot showing the log_2_fold change in metabolite abundance versus -log_10_(q value) from *FLVCR1*-knockout HEK293T and HeLa cells expressing a vector control or *FLVCR1* cDNA cultured in choline replete media for 24 hours. The dotted line indicates the significance threshold of FDR q<1% (-log_10_(q value) = 2). Statistical significance was determined by multiple two-tailed unpaired *t*-tests with discovery determined using the Two-stage linear step-up procedure of Benjamini, Krieger and Yekutieli. **c)** Protein normalized total phosphocholine, betaine, and glycerophosphocholine abundance after incubation with [1,2-^13^C_2_]Choline for 24 hours in *FLVCR1*-knockout HeLa cells expressing a control vector or *FLVCR1* cDNA. Bars represent mean ± standard deviation; n = 3. Statistical significance determined by two-tailed unpaired *t*-test and displayed p-values compare M+2 metabolite abundance between *FLVCR1* KO and *FLVCR1* KO +*FLVCR1* cDNA. **d)** Fractional labeling of phosphocholine, betaine, and glycerophosphocholine after incubation with [1,2-^13^C_2_]Choline for the indicated timepoints in *FLVCR1*-knockout HEK293T cells expressing a control vector or *FLVCR1* cDNA. Bars represent mean ± standard deviation; n = 3. **e)** Fractional labeling of phosphocholine, betaine, and glycerophosphocholine after incubation with [1,2-^13^C_2_]Choline for 24 hours *FLVCR1*-knockout HeLa cells expressing a control vector or *FLVCR1* cDNA. Bars represent mean ± standard deviation; n = 3. **f)** Lidipomics of *FLVCR1*-knockout HEK293T cells expressing a control vector or *FLVCR1* cDNA cultured in choline replete media for 24 hours. Dot plot of individual lipid species of *FLVCR1*-knockout cells relative to *FLVCR1* KO +*FLVCR1* cDNA cells and grouped by lipid class – phosphatidylcholine (PC), phosphatidylethanolamine (PE), phosphatidylserine (PS), and triglyceride (TG). Dotted line represents lipid species abundance of *FLVCR1* KO +*FLVCR1* cDNA cells. Median log_2_ fold change of *FLVCR1* KO cells denoted by black line for each lipid class; n = 3. **g)** Quantification of LipidTOX staining of *FLVCR1*-knockout HEK293T cells expressing a control vector or *FLVCR1* cDNA. Violin plot showing quantification of maximum fluorescence intensity per cell. Statistical significance was determined by two-tailed unpaired *t*-test. n = 28 cells *FLVCR1* KO; n = 39 cells *FLVCR1* KO +*FLVCR1* cDNA **h)** Metabolite levels normalized by endogenous leucine between *Flvcr1*^KO^(n = 4) and *Flvcr1*^HET^ (n = 6) plus *Flvcr1*^WT^ (n = 2) mice. Bars represent mean ± standard deviation. Statistical significance determined by two-tailed unpaired *t*-test.. **i)** Immunoblotting of indicated proteins in HEK293T cells treated with 20μM Hemin or vehicle control for 24 hours. **j)** Protein normalized total phosphocholine and betaine abundance after incubation with [1,2-^13^C_2_]Choline for 3 hours in HEK293T cells treated with 20μM Hemin or vehicle control for 24 hours. Bars represent mean ± standard deviation; n = 3. Statistical significance determined by two-tailed unpaired *t*-test and displayed p-values compare M+2 metabolite abundance between vehicle and hemin treated cells. **k)** Immunoblotting of indicated proteins in *CHKA*-knockout HEK293T cells expressing a vector control or *CHKA* cDNA. **l)** Protein normalized total phosphocholine, betaine, and glycerophosphocholine abundance after incubation with [1,2-^13^C_2_]Choline for 24 hours in *CHKA*-knockout HEK293T cells expressing a control vector or *CHKA* cDNA. Bars represent mean ± standard deviation; n = 3. Statistical significance determined by two-tailed unpaired *t*-test and displayed p-values compare M+2 metabolite abundance between *CHKA* KO and *CHKA* KO +*CHKA* cDNA. **m)** Immunoblotting of indicated proteins in *FLVCR1*-knockout HEK293T and HeLa cells expressing a vector control or *FLVCR1* cDNA.

**Supplemental Figure 3. FLVCR1 mediated choline uptake is hemicholinium-3 and Na+ independent. a)** Bulk-RNA expression of SLC5A7 and FLVCR1 across human tissues from Human Protein Atlas HPA Consensus dataset (derived from HPA RNAseq and Genotype-Tissue Expression GTEx project RNAseq). Tissues are color-coded by organ or organ system **b)** Uptake of [Methyl-^3^H]-Choline in *FLVCR1*-knockout HeLa cells expressing a vector control or *FLVCR1* cDNA incubated for indicated timepoint. Data represented as mean ± standard deviation and normalized by seeded cell number; n = 3. Statistical significance was determined by multiple two-sided unpaired *t*-tests with correction for multiple comparisons using the Holm-Sidak method. t=1 minute p=7×10^−6^; t =2 minutes p<1×10^−6^, t=5 minutes p=1.1×10^−6^, t=15 minutes p=2.4×10^−5^, and 30 minutes p=4×10^−6^. **c)** Uptake of [Methyl-^3^H]-Choline in *FLVCR1*-knockout HeLa cells expressing a vector control or *FLVCR1* cDNA incubated with indicated dose for 30 minutes. Data represented as mean ± standard deviation and normalized by seeded cell number; n = 3. Statistical significance was determined by multiple two-sided unpaired *t*-tests with correction for multiple comparisons using the Holm-Sidak method. 1.34μM, 2.69μM, 5.38μM, 10.75μM, and 21.5μM p<1×10^−6^. **d)** Protein normalized phosphocholine abundance in *FLVCR1*-knockout HEK293T cells expressing a vector control or *FLVCR1* cDNA after incubation with 1mM phosphocholine or 1mM choline for 24 hours. Bars represent mean ± standard deviation; n = 3. Statistical significance determined by two-tailed unpaired *t*-test. **e)** Uptake of 5.38 μM [Methyl-^3^H]-Choline in *FLVCR1*-knockout HEK293T cells expressing a vector control or *FLVCR1* cDNA incubated with indicated dose of hemicholinium-3 for 30 minutes. Data represented as mean ± standard deviation and normalized by seeded cell number; n = 3. Statistical significance was determined by two-sided unpaired *t*-test. Within HEK293T cells expressing a vector control or *FLVCR1* cDNA 0μM hemicholinium-3 was compared to other doses. 0.01μM hemicholinium-3: *FLVCR1* KO p=0.416, *FLVCR1* KO +*FLVCR1* cDNA p=0.053; 0.1μM *FLVCR1* KO p=0.037, *FLVCR1* KO +*FLVCR1* cDNA p=0.011; 1μM *FLVCR1* KO p=8.7×10^−3^, *FLVCR1* KO +*FLVCR1* cDNA p=2.7×10^−3^; 10μM *FLVCR1* KO p=1×10^−3^, *FLVCR1* KO +*FLVCR1* cDNA p=1×10^−3^. **f)** Uptake of [Methyl-^3^H]-Choline in *FLVCR1*-knockout HEK293T cells expressing a vector control or *FLVCR1* cDNA incubated with indicated dose of hemicholinium-3 for 30 minutes relative to uptake at 0*uM* of hemicholinium-3. Statistical significance was determined by multiple two-sided unpaired *t*-tests with correction for multiple comparisons using the Holm-Sidak method. 0.01μM p=0.159; 0.1μM p=0.139; 1μM p=0.197; 10μM p=0.118. **g)** Uptake of 5.38μM [Methyl-^3^H]-Choline in *FLVCR1*-knockout HeLa cells expressing a vector control or *FLVCR1* cDNA incubated with indicated dose of hemicholinium-3 for 30 minutes. Data represented as mean ± standard deviation and normalized by seeded cell number; n = 3. Statistical significance was determined by two-sided unpaired *t*-test. Within HeLa cells expressing a vector control or *FLVCR1* cDNA 0μM hemicholinium-3 was compared to other doses. 0.01μM hemicholinium-3: *FLVCR1* KO p=0.061, *FLVCR1* KO +*FLVCR1* cDNA p=0.023; 0.1μM *FLVCR1* KO p=0.346, *FLVCR1* KO +*FLVCR1* cDNA p=7×10^−3^; 1μM *FLVCR1* KO p=4×10^−3^, *FLVCR1* KO +*FLVCR1* cDNA p=3×10^−4^; 10μM *FLVCR1* KO p=9×10^−4^, *FLVCR1* KO +*FLVCR1* cDNA p=2×10^−4^. **h)** Uptake of [Methyl-^3^H]-Choline in *FLVCR1*-knockout HEK293T cells expressing a vector control or *FLVCR1* cDNA incubated with indicated dose of hemicholinium-3 for 30 minutes relative to uptake at 0*uM* of hemicholinium-3. Statistical significance was determined by multiple two-sided unpaired *t*-tests with correction for multiple comparisons using the Holm-Sidak method. 0.01μM p=7.2×10^−5^; 0.1μM p=9.9×10^−3^; 1μM p=0.366; 10μM p=0.332. **i)** Uptake of [Methyl-^3^H]-Choline in *FLVCR1*-knockout HEK293T cells expressing an empty vector (EV) control, *FLVCR1*, or *SLC5A7* cDNA for 30 minutes in Na^+^ or K^+^ Krebs-Ringer buffer. Data represented as mean ± standard deviation and normalized by seeded cell number; n = 3. Statistical significance determined by two-tailed unpaired *t*-test. **j)** Cumulative log2 fold change in cell number after continuous culture in choline depleted or replete media of *FLVCR1*-knockout HEK293T and HeLa cells expressing a vector control or *FLVCR1* cDNA. **k)** Log_2_ fold change in cell number of *FLVCR1*-knockout HeLa and HEK293T cells expressing a vector control or *FLVCR1* cDNA grown in choline depleted or choline replete media supplemented with 1% or 10% dialyzed FBS (dFBS). Bars represent mean ± standard deviation; n = 6. Statistical significance determined by two-tailed unpaired *t*-test.

**Supplemental Figure 4. FLVCR2, the mammalian paralog of FLVCR1, can compensate for FLVCR1 loss. a)** CRISPR gene scores in *FLVCR1*-knockout HEK293T cells expressing a vector control grown in standard conditions (*y*-axis) or supplemented with 1mM choline (*x*-axis). Top scoring hits color coded and previously reported potential choline transporters highlighted in gray. Pearson correlation coefficient, two-sided. **b)** Top 15 scoring genes differentially required for *FLVCR1*-knockout HEK293T cells compared to cDNA addback (left) or 1mM Choline supplementation (right). **c)** Barcode plot of gene scores across genetic screens. Left = *FLVCR1* KO – *FLVCR1* KO +*FLVCR1* cDNA; Center: *FLVCR1* KO – *FLVCR1* KO +1mM Choline; Right = *FLVCR1* KO +*FLVCR1* cDNA – *FLVCR1* KO +1mM Choline. **d)** Lipidomic analysis of *FLVCR1*-knockout or *FLVCR1* cDNA complemented HEK293T cells expressing an empty vector control (pLV2 EV) or two different sgRNA targeting *SPNS1*. Violin plot of individual lysophosphatidylcholine species; n = 3. **e)** Phylogenetic tree of FLVCR1 homologs across model organisms (derived from TreeFam). **f)** Bulk-RNA expression of FLVCR2 across human tissues from Human Protein Atlas HPA Consensus dataset (derived from HPA RNAseq and Genotype-Tissue Expression GTEx project RNAseq). Tissues are color-coded by organ or organ system. **g)** Uptake of 21.5 μM [Methyl-^3^H]-Choline in *FLVCR1*-knockout HEK293T cells expressing an empty vector (EV) control, *SLC44A1* cDNA, *FLVCR2* cDNA, *FLVCR1* cDNA, or *SLC5A7* cDNA for 30 minutes. Data represented as mean ± standard deviation and normalized by seeded cell number; n = 3. Statistical significance determined by two-tailed unpaired *t*-test

**Supplemental Figure 5. Choline deprivation in *FLVCR1*-knockout cells rewires the mitochondrial lipidome and proteome. a)** Immunoblotting of indicated proteins in *FLVCR1*-knockout HeLa cells expressing a vector control or *FLVCR1* cDNA cultured in choline depleted or choline replete media supplemented with 1% or 10% dialyzed FBS (dFBS). **b)** Immunoblotting of indicated proteins in *FLVCR1*-knockout HEK293T cells expressing a vector control or *FLVCR1* cDNA cultured in choline depleted media supplemented with 1% or 10% dFBS. **c)** Immunoblotting of *CHKA*-knockdown or empty vector control HeLa cells cultured in choline depleted media supplemented with 1% or 10% dFBS. **d)** Immunoblotting of HeLa cells pretreated with varying amounts of phosphocholine for 24hrs followed by treatment with CHKA inhibitor RSM932A (1μM) or vehicle for 24 hours with supplemented phosphocholine. **e)** Immunoblotting of indicated proteins in HEK293T cells expressing an empty vector control (pLV2 EV) or two different sgRNA targeting DELE1 and treated with 1μM oligomycin or vehicle for 24 hours. **f)** Immunoblotting of indicated proteins in HEK293T cells expressing an empty vector control (pLV2 EV) or two different sgRNA targeting DELE1 and treated with 0.5μg/mL tunicamycin or vehicle for 24 hours. **g)** Immunoblotting of indicated proteins of input or immunoprecipitation (IP) samples of *FLVCR1*-knockout HeLa cells expressing a vector control or *FLVCR1* cDNA with OMP25 Myc or HA tags following culture in choline depleted or replete media. **h)** PCA of MitoIP proteomics *FLVCR1*-knockout HeLa cells expressing a vector control or *FLVCR1* cDNA cultured in choline depleted or choline replete media. **i)** Volcano plot showing the log_2_ fold change in protein abundance versus -log_10_(q value) from the MitoIP of *FLVCR1*-knockout HeLa cells expressing a vector control or *FLVCR1* cDNA cultured in choline depleted media. The dotted line indicates the significance threshold of FDR q<1% (-log_10_(q value) = 2). Proteins from a consensus multi-drug signature of mitochondrial stress are indicated in red. Gene set enrichment analysis (GSEA) was used to determine statistically significant enrichment of the Quiros gene set in the proteomics data.

**Table 1. Full list of predicted metabolite-plasma membrane transporter associations from METSIM analysis.**

## METHODS

### Cell Lines and Growth Medium

HeLa and HEK293T cells were obtained from the ATCC. Cell line identity was authenticated using STR profiling and cell lines were verified to be free of mycoplasma contamination. Under standard culture conditions, cell lines were grown in RPMI 1640 medium (Gibco #11875-093) containing 2mM glutamine supplemented with 10% fetal bovine serum (FBS) (Sigma-Aldrich #12306C) and 1% penicillin streptomycin (Gibco #15140-122). All cells were maintained at 37°C, 21% O_2_, and 5% CO_2._Choline depleted and replete medium was made from powdered RPMI 1650 Medium w/o L-Glutamine, L-Methionine, Choline Chloride, Folic Acid and Vitamin B12 (US Biological Life Sciences #R8999-21). Once reconstituted, missing metabolites were added at the concentration found in standard RPMI 1640 medium: 2mM L-glutamine (Alfa Aesar #J60573), 100.7μM L-methionine (Alfa Aesar #J61904), 21.4μM choline chloride (MP Biomedicals #101386), 2.3μM folic acid (Alfa Aesar #J62937), and 3.7nM vitamin B12 (Alfa Aesar #A14894). Choline depleted and replete medium was supplemented with 10% dialyzed fetal bovine serum (dFBS) (Gibco #26400-044) and 1% penicillin streptomycin (Gibco #15140-122), unless otherwise noted.

### Chemicals and Compounds

The following chemicals and compounds were used with doses enumerated in the figures and figure legends: Choline chloride (MP Biomedicals #67-48-1), Phosphocholine chloride calcium salt tetrahydrate (Sigma Aldrich #P0378), RSM932A (Cayman Chemical #215518), Tunicamycin (Tocris #3516), Oligomycin (Millipore #495455), Hemin (Sigma Aldrich #H9039), Hemicholinium-3 (Sigma Aldrich #H108). For all treatments an equal amount of respective vehicle was used as the untreated control.

### Generation of Knockout and Overexpression Constructs and Cell Lines

sgRNAs (listed in ‘Oligonucleotide sequences’) were synthesized by IDT and cloned into lentiCRISPR-v2 puro (Addgene #982990) or lentiCRISPR-v2 GFP (Addgene #75159) linearized by BsmBI using a T4 ligase (NEB #M0202). cDNAs (listed in ‘Oligonucleotide sequences’) were synthesized by IDT or Twist Biosciences and cloned into pMXS-IRES-BLAST (Cell Biolabs #RTV-016) linearized with BamHI and NotI using Gibson Assembly (NEB #E2611). All construct sequences were validated by Sanger Sequencing. sgRNA expressing vectors along with lentiviral packaging vectors Delta-VPR and VSV-G were transfected into HEK293T cells using X-tremeGENE 9 DNA Transfection reagent (Roche #6364787001). cDNA expressing vectors along with retroviral packaging vectors Gag-Pol and VSG-G were transfected into HEK293T cells using X-tremeGENE 9 DNA Transfection reagent (Roche #6364787001). Virus-containing supernatant was collected 48 hours after transfected and passed through a 0.45μm filter. Target cells in 6-well tissue culture plates were spin-infected with virus and 4μg/mL polybrene by centrifugation at 2,200rpm for 80 minutes. Cells were selected by puromycin (lentiCRISPR-v2 puro), blasticidin (pMXS-IRES-BLAST), or FACS (lentiCRISPR-v1 GFP). Transduced cells were subsequently single-cell cloned. For all constructs generated the matching vector without insert was used as a control.

### Oligonucleotide Sequences

#### sgRNAs

Human *FLVCR1*_sg5 F: 5’-GTAGGCCAGCATGTACACCA-3’

Human *FLVCR1*_sg5 R: 5’-TGGTGTACATGCTGGCCTAC-3’

Human *CHKA*_sg2 F: 5’-GAAAACCAAATTCTGCACCG-3’

Human *CHKA*_sg2 R: 5’-CGGTGCAGAATTTGGTTTTC-3’

Human *KIAA0141/DELE1*_sg1 F: 5’-GAGCAGTGTTCTACCCGCTG-3’

Human *KIAA0141/DELE1*_sg1 R: 5’-CAGCGGGTAGAACACTGCTC-3’

Human *KIAA0141/DELE1*_sg2 F: 5’-GGAACAGAGAACATGAAGAG-3’

Human *KIAA0141/DELE1*_sg2 R: 5’-CTCTTCATGTTCTCTGTTCC-3’

Human *EIF2AK3/PERK*_sgRNA F: 5’-GCAGAGGCCGGGCTGAGACG-3’

Human *EIF2AK3/PERK*_sgRNA R: 5’-CGTCTCAGCCCGGCCTCTGC-3’

Human *EIF2AK2/PKR*_sgRNA F: 5’-GGCGGCGGCGCAGGTGAGCA-3’

Human *EIF2AK2/PKR*_sgRNA R: 5’-TGCTCACCTGCGCCGCCGCC-3’ Human *GCN2_sgRNA* F: 5’-GAACTGGCCAAGAAACACTG-3’

Human *GCN2_sgRNA* R: 5’-CAGTGTTTCTTGGCCAGTTC-3’

Human *EIF2AK1/HRI*_sgRNA F: 5’-CGCGAAGAGGAGGGCGACG-3’

Human *EIF2AK1/ HRI*_sgRNA R: 5’-CGTCGCCCTCCTCTTCGCGc-3’

Human *SPNS1*_sg1 F: 5’- gCTACATGGACCGCTTCACCG-3’

Human *SPNS1*_sg1 R: 5’- CGGTGAAGCGGTCCATGTAGc-3’

Human *SPNS1*_sg2 F: 5’- gAAGTGAGTCCCTACCAGCCA-3’

Human *SPNS1*_sg2 R: 5’-TGGCTGGTAGGGACTCACTTc-3’

#### cDNAs (Codon Optimized; 5’-3’)

##### Human *FLVCR1*

ATGGCCCGCCCCGATGACGAAGAGGGAGCTGCCGTAGCTCCTGGCCATCCACTGGCCAAGGGGT ATCTGCCTCTCCCACGGGGTGCCCCTGTGGGAAAAGAATCCGTCGAACTTCAAAATGGCCCTAAG GCCGGAACTTTTCCTGTTAACGGCGCGCCTCGTGATTCTCTGGCAGCTGCTAGCGGTGTATTGGG TGGACCACAAACCCCACTTGCACCCGAAGAAGAAACACAAGCCAGATTGCTGCCAGCCGGCGCG GGCGCCGAAACCCCCGGCGCCGAATCATCACCACTTCCTCTGACAGCCCTGTCTCCTAGACGATT TGTCGTCCTGCTCATATTTAGTCTTTACTCTCTCGTGAACGCATTCCAATGGATACAGTATTCAATA ATCTCAAATGTGTTTGAAGGATTTTATGGCGTGACTCTGTTGCATATAGATTGGCTTAGCATGGTCT ATATGCTCGCTTATGTTCCTCTGATATTTCCAGCTACATGGCTTCTCGATACTAGGGGTCTCCGCC TTACAGCACTCCTCGGGTCAGGTCTGAATTGTCTCGGAGCATGGATAAAATGTGGATCCGTCCAA CAACACCTGTTTTGGGTGACGATGCTCGGGCAATGTCTTTGTTCAGTTGCACAAGTCTTTATTCTC GGGCTGCCATCAAGAATTGCTTCTGTCTGGTTCGGCCCTAAGGAAGTAAGCACGGCCTGCGCAAC GGCAGTCCTTGGGAACCAACTTGGCACCGCTGTCGGGTTCCTGCTGCCACCCGTGCTGGTGCCT AATACCCAAAACGATACGAACCTGCTCGCCTGCAACATTTCCACTATGTTCTACGGCACCTCTGCA GTGGCGACTCTCCTGTTCATACTGACCGCGATCGCGTTCAAGGAGAAGCCGCGCTACCCACCGTC ACAAGCCCAGGCTGCCTTGCAGGATTCACCACCCGAGGAATATAGCTACAAGAAGAGCATCAGGA ATTTGTTCAAGAATATCCCGTTCGTGCTGTTGCTGATTACCTACGGCATTATGACAGGGGCATTTTA TAGTGTGTCAACTCTCCTCAATCAAATGATTCTCACATATTATGAAGGTGAGGAAGTGAACGCAGG CAGAATCGGCCTGACCCTTGTGGTCGCCGGCATGGTAGGTAGCATCCTGTGCGGTTTGTGGCTC GACTACACCAAGACGTATAAGCAAACGACCCTTATTGTCTACATACTGTCCTTCATCGGCATGGTC ATTTTCACCTTTACTCTGGATCTGAGGTACATCATTATTGTCTTCGTGACCGGTGGCGTCCTGGGG TTCTTTATGACAGGCTATCTGCCCCTGGGGTTCGAGTTCGCCGTCGAGATTACATATCCCGAGTCA GAGGGGACATCTTCCGGGCTGCTGAACGCAAGCGCCCAAATTTTCGGTATCCTGTTTACCCTGGC CCAGGGGAAATTGACCTCTGATTACGGACCGAAAGCTGGTAATATCTTCCTTTGCGTGTGGATGTT CATTGGGATTATCCTGACGGCCCTGATTAAATCCGACCTTCGGCGGCATAATATCAACATTGGGAT CACCAACGTCGACGTCAAGGCCATCCCCGCCGATAGCCCGACTGATCAGGAGCCTAAGACTGTG ATGCTGAGTAAACAATCTGAGAGTGCTATCTAG

##### Human *FLVCR2*

ATGGTAAACGAGGGACCGAATCAAGAGGAATCTGACGATACACCAGTACCCGAATCAGCGTTGCAGGCCGATCCGTCAGTTTCTGTGCACCCTTCCGTGTCAGTTCACCCATCCGTTTCTATTAATCCGTCCGTGAGCGTACATCCGTCATCCTCTGCTCATCCTTCTGCACTCGCTCAGCCTTCAGGACTTGCCCATCCATCATCTTCAGGGCCCGAAGATCTGAGTGTCATAAAAGTCTCCCGGAGAAGGTGGGCAGTTGTTTTAGTCTTCTCTTGTTATAGCATGTGCAATTCTTTCCAATGGATCCAGTACGGGTCTATTAACAACATATTTATGCATTTCTATGGAGTGTCTGCTTTCGCTATCGATTGGCTCTCAATGTGTTATATGCTCACCTATATACCGCTTTTGTTACCGGTCGCCTGGTTGCTTGAGAAATTTGGGTTGAGAACAATCGCGCTGACCGGTAGCGCACTGAATTGTCTTGGTGCTTGGGTTAAATTGGGGTCTTTGAAACCCCACCTGTTCCCTGTGACAGTTGTTGGTCAACTGATTTGTAGTGTTGCTCAAGTCTTTATTCTTGGAATGCCAAGCAGAATAGCAAGTGTATGGTTTGGCGCAAACGAAGTCTCCACCGCTTGTAGCGTAGCCGTGTTCGGGAACCAATTAGGCATAGCCATAGGCTTTCTGGTGCCGCCAGTACTTGTGCCAAATATCGAGGATCGAGATGAACTGGCATATCATATTTCTATAATGTTTTACATCATTGGTGGCGTTGCTACCTTATTGCTGATTCTCGTGATTATCGTGTTTAAAGAAAAGCCAAAGTACCCGCCTTCAAGAGCTCAGAGCCTGTCTTACGCGCTGACGAGCCCGGACGCTTCTTATCTCGGCTCAATTGCTAGACTTTTCAAGAACCTTAATTTCGTCTTGCTGGTGATTACATACGGGCTCAACGCAGGCGCATTCTACGCGCTGAGCACCCTGTTGAACCGGATGGTAATTTGGCATTATCCCGGCGAAGAGGTTAACGCCGGGCGCATAGGTTTAACCATTGTGATCGCCGGCATGCTGGGAGCGGTCATAAGCGGTATTTGGTTAGACAGATCAAAGACTTACAAGGAAACTACTTTAGTGGTGTACATTATGACCTTGGTCGGAATGGTTGTTTATACATTCACACTGAATTTGGGCCATCTTTGGGTGGTTTTTATTACCGCTGGAACCATGGGTTTCTTCATGACAGGATACTTGCCCTTGGGCTTCGAATTCGCAGTTGAATTAACCTATCCTGAGTCCGAGGGGATAAGCTCTGGACTTCTGAATATCTCCGCTCAAGTGTTCGGCATTATTTTCACGATTAGCCAAGGACAAATCATAGATAATTACGGGACAAAACCAGGCAATATTTTCTTATGCGTCTTTCTCACATTAGGCGCTGCTCTGACAGCTTTTATCAAAGCTGACTTGCGCCGCCAAAAGGCTAACAAGGAGACCCTCGAAAATAAGCTGCAGGAAGAAGAAGAAGAATCAAATACGTCCAAGGTACCGACCGCCGTCTCCGAAGACCACTTGTA G

##### Human *SLC44A1*

ATGGGCTGCTGCAGCTCCGCCTCCTCCGCCGCGCAGAGCTCtAAgCGAGAgTGGAAaCCaCTGGAGGACCGTAGCTGCACAGACATACCATGGCTGCTGCTCTTCATCCTCTTCTGCATTGGGATGGGATTTATTTGTGGCTTTTCAATAGCAACAGGTGCAGCAGCAAGACTAGTGTCAGGATACGACAGCTATGGAAATATCTGTGGGCAGAAAAATACAAAGTTGGAAGCAATACCAAACAGcGGtATGGAtCAtACgCAaCGGAAGTATGTATTCTTTTTGGATCCATGCAACCTGGACTTGATAAACCGGAAGATTAAGTCTGTAGCACTGTGTGTAGCAGCGTGTCCAAGGCAAGAACTGAAAACTCTGAGTGATGTTCAGAAGTTTGCAGAGATAAATGGTTCAGCCCTATGTAGCTACAACCTAAAGCCTTCTGAATACACTACATCTCCAAAATCTTCTGTTCTCTGCCCCAAACTACCAGTTCCAGCGAGTGCACCTATTCCATTCTTCCATCGCTGTGCTCCTGTGAACATTTCCTGCTATGCCAAGTTTGCAGAGGCCCTGATCACCTTTGTCAGTGACAATAGTGTCTTACACAGGCTGATTAGTGGAGTAATGACCAGCAAAGAAATTATATTGGGACTTTGCTTGTTATCACTAGTTCTATCCATGATTTTGATGGTGATAATCAGGTATATATCAAGAGTACTTGTGTGGATCTTAACGATTCTGGTCATACTCGGTTCACTTGGAGGCACAGGTGTACTATGGTGGCTGTATGCAAAGCAAAGAAGGTCTCCCAAAGAAACTGTTACTCCTGAGCAGCTTCAGATAGCTGAAGACAATCTTCGGGCCCTCCTCATTTATGCCATTTCAGCTACAGTGTTCACAGTGATCTTATTCCTGATAATGTTGGTTATGCGCAAACGTGTTGCTCTTACCATCGCCTTGTTCCACGTAGCTGGCAAGGTCTTCATTCACTTGCCACTGCTAGTCTTCCAACCCTTCTGGACTTTCTTTGCTCTTGTCTTGTTTTGGGTGTACTGGATCATGACACTTCTTTTTCTTGGCACTACCGGCAGTCCTGTTCAGAATGAGCAAGGCTTTGTGGAGTTCAAAATTTCTGGGCCTCTGCAGTACATGTGGTGGTACCATGTGGTGGGCCTGATTTGGATCAGTGAATTTATTCTAGCATGTCAGCAGATGACAGTGGCAGGAGCTGTGGTAACATACTATTTTACTAGGGATAAAAGGAATTTGCCATTTACACCTATTTTGGCATCAGTAAATCGCCTTATTCGTTACCACCTAGGTACGGTGGCAAAAGGATCTTTCATTATCACATTAGTCAAAATTCCGCGAATGATCCTTATGTATATTCACAGTCAGCTCAAAGGAAAGGAAAATGCTTGTGCACGATGTGTGCTGAAATCTTGCATTTGTTGCC TTTGGTGTCTTGAAAAGTGCCTAAATTATTTAAATCAGAATGCATACACAGCCACAGCTATCAACAGCACCAACTTCTGCACCTCAGCAAAGGATGCCTTTGTCATTCTGGTGGAGAATGCTTTGCGAGTGGCTACCATCAACACAGTAGGAGATTTTATGTTATTCCTTGGCAAGGTGCTGATAGTCTGCAGCACAGGTTTAGCTGGGATTATGCTGCTCAACTACCAGCAGGACTACACAGTATGGGTGCTGCCTCTGATCATCGTCTGCCTCTTTGCTTTCCTAGTCGCTCATTGCTTCCTGTCTATTTATGAAATGGTAGTGGATGTATTATTCTTGTGTTTTGCCATTGATACAAAATACAATGATGGGAGCCCTGGCAGAGAATTCTATATGGATAAAGTGCTGATGGAGTTTGTGGAAAACAGTAGGAAAGCAATGAAAGAAGCTGGTAAGGGAGGCGTCGCTGATTCCAGAGAGCTAAAGCCGATGGCTTCGGGAGCAAGTTCTGCTTGA

##### Human *SLC5A7*

ATGGCTTTCCATGTGGAAGGACTGATAGCTATCATCGTGTTCTACCTTCTAATTTTGCTGGTTGGAATATGGGCTGCCTGGAGAACCAAAAACAGTGGCAGCGCAGAAGAGCGCAGCGAAGCCATCATAGTTGGTGGCCGAGATATTGGTTTATTGGTTGGTGGATTTACCATGACAGCTACCTGGGTCGGAGGAGGGTATATCAATGGCACAGCTGAAGCAGTTTATGTACCAGGTTATGGCCTAGCTTGGGCTCAGGCACCAATTGGATATTCTCTTAGTCTGATTTTAGGTGGCCTGTTCTTTGCAAAACCTATGCGTTCAAAGGGGTATGTGACCATGTTAGACCCGTTTCAGCAAATCTATGGAAAACGCATGGGCGGACTCCTGTTTATTCCTGCACTGATGGGAGAAATGTTCTGGGCTGCAGCAATTTTCTCTGCTTTGGGAGCCACCATCAGCGTGATCATCGATGTGGATATGCACATTTCTGTCATCATCTCTGCACTCATTGCCACTCTGTACACACTGGTGGGAGGGCTCTATTCTGTGGCCTACACTGATGTCGTTCAGCTCTTTTGCATTTTTGTAGGGCTGTGGATCAGCGTCCCCTTTGCATTGTCACATCCTGCAGTCGCAGACATCGGGTTCACTGCTGTGCATGCCAAATACCAAAAGCCGTGGCTGGGAACTGTTGACTCATCTGAAGTCTACTCTTGGCTTGATAGTTTTCTGTTGTTGATGCTGGGTGGAATCCCATGGCAAGCATACTTTCAGAGGGTTCTCTCTTCTTCCTCAGCCACCTATGCTCAAGTGCTGTCCTTCCTGGCAGCTTTCGGGTGCCTGGTGATGGCCATCCCAGCCATACTCATTGGGGCCATTGGAGCATCAACAGACTGGAACCAGACTGCATATGGGCTTCCAGATCCCAAGACTACAGAAGAGGCAGACATGATTTTACCAATTGTTCTGCAGTATCTCTGCCCTGTGTATATTTCTTTCTTTGGTCTTGGTGCAGTTTCTGCTGCTGTTATGTCATCAGCAGATTCTTCCATCTTGTCAGCAAGTTCCATGTTTGCACGGAACATCTACCAGCTTTCCTTCAGACAAAATGCTTCGGACAAAGAAATCGTTTGGGTTATGCGAATCACAGTGTTTGTGTTTGGAGCATCTGCAACAGCCATGGCCTTGCTGACGAAAACTGTGTATGGGCTCTGGTACCTCAGTTCTGACCTTGTTTACATCGTTATCTTCCCCCAGCTGCTTTGTGTACTCTTTGTTAAGGGAACCAACACCTATGGGGCCGTGGCAGGTTATGTTTCTGGCCTCTTCCTGAGAATAACTGGAGGGGAGCCATATCTGTATCTTCAGCCCTTGATCTTCTACCCTGGCTATTACCCTGATGATAATGGTATATATAATCAGAAATTTCCATTTAAAACACTTGCCATGGTTACATCATTCTTAACCAACATTTGCATCTCCTATCTAGCCAAGTATCTATTTGAAAGTGGAACCTTGCCACCTAAATTAGATGTATTTGATGCTGTTGTTGCAAGACACAGTGAAGAAAACATGGATAAGACAATTCTTGTCAAAAATGAAAATATTAAATTAGATGAACTTGCACTTGTGAAGCCACGACAGAGCATGACCCTCAGCTCAACTTTCACCAATAAAGAGGCCTTCCTTGATGTTGATTCCAGTCCAGAAGGGTCTGGGACTGAAGATAATTTACAGTGA

##### Human *CHKA*

ATGAAGACAAAGTTTTGTACAGGTGGGGAAGCAGAACCAAGTCCCCTGGGCCTCCTCCTCAGTTGCGGATCAGGATCCGCTGCACCTGCCCCAGGTGTCGGACAACAAAGAGATGCAGCATCAGATTTGGAAAGCAAACAACTTGGTGGTCAGCAACCACCACTTGCTCTTCCCCCGCCCCCACCACTCCCTTTGCCACTTCCACTCCCTCAACCACCACCCCCACAACCACCTGCGGATGAACAACCCGAACCGAGAACTCGAAGGCGTGCTTACCTTTGGTGTAAAGAATTTCTTCCAGGAGCATGGCGTGGGCTTCGGGAAGATGAATTTCATATTTCTGTGATACGTGGCGGACTCTCTAATATGTTGTTTCAATGTAGCCTTCCAGATACAACGGCAACTCTGGGAGACGAACCCCGTAAGGTTCTGCTCCGCCTTTACGGCGCTATTCTCCAAATGCGGAGCTGCAACAAGGAAGGGAGCGAGCAGGCGCAAAAGGAGAACGAGTTCCAGGGAGCAGAAGCAATGGTGTTGGAATCCGTCATGTTCGCAATCCTGGCGGAACGTAGTCTGGGCCCCAAGCTGTACGGGATTTTCCCACAGGGAAGATTGGAACAATTTATACCATCTAGGCGCCTCGACACAGAGGAGCTGTCCCTGCCTGACATCAGCGCTGAGATAGCTGAAAAGATGGCAACCTTCCACGGAATGAAGATGCCCTTTAACAAAGAGCCGAAGTGGCTGTTCGGAACGATGGAGAAATACCTGAAAGAGGT ATTGCGGATCAAGTTCACCGAAGAGAGCCGCATCAAGAAACTGCATAAGCTCCTTTCCTATAACCTTCCTCTGGAGTTGGAGAATCTTAGGAGTCTGCTGGAGAGCACCCCGTCCCCCGTAGTCTTCTGCCACAACGATTGCCAAGAGGGCAACATACTGCTGCTCGAGGGAAGGGAAAACAGCGAGAAGCAAAAGCTTATGCTGATCGACTTTGAGTATTCTTCTTATAACTATCGCGGGTTTGATATAGGCAACCACTTTTGCGAATGGATGTACGACTACAGTTACGAGAAGTATCCCTTCTTTAGGGCGAATATACGCAAATACCCAACAAAGAAGCAGCAATTGCACTTCATCTCAAGCTATCTGCCAGCCTTTCAGAACGATTTCGAGAATCTGTCCACCGAGGAGAAGTCTATCATCAAGGAAGAGATGCTTCTCGAGGTCAACCGGTTCGCTTTGGCTAGTCACTTTCTTTGGGGCCTTTGGTCAATCGTTCAGGCTAAAATCTCCAGCATCGAGTTCGGTTATATGGATTATGCTCAGGCTCGATTCGACGCATACTTTCATCAAAAGCGCAAACTCGGCGT TTAA

### Immunoblotting

Cells were lysed in either RIPA buffer (10mM Tris-HCl pH 7.5, 150mM NaCl, 1mM EDTA, 1% Triton X- 100, 0.1% SDS) or transmembrane buffer (10mM Tris-HCl pH 7.4, 1mM EDTA, 1% Triton X-100, 2% SDS, 0.1% CHAPS) supplemented with protease inhibitors (EMD Millipore #535140 or Sigma-Aldrich #11836170001) and phosphatase inhibitors (Roche #490685001). For mitochondrial immunoprecipitation experiments, samples were lysed in 1% Triton X-100 Buffer (50mM Tris-HCl pH 7.4, 150mM NaCl, 1mM EDTA, 1% Triton X-100) supplemented with protease inhibitors (Sigma-Aldrich #11836170001). Lysates were sonicated, centrifuged at 14,000 rpm, and supernatant collected. Total protein was quantified using BCA Protein Assay Kit (Thermo Fisher #23227) with provided albumin standard used as a protein standard. Samples were resolved on 12%, 4-12%, or 10-20% Tris-Glycine gels (Invitrogen #XP00125BOX, #XP04125BOX, #XP10205BOX) and analyzed by standard immunoblotting techniques. Briefly, gels were transferred in CAPS buffer (10mM CAPS, 10% ethanol) to PVDF membranes (EMD Millipore #1PVH00010) and incubated with primary antibodies at 4°C overnight (ATF4, CST #11815S; CHOP, CST #2895S; GAPDH, Genetex #GTX627408; Calreticulin, CST #12238P; LAMP1, CST #9091PP; COXIV, CST #4850S; Beta Tubulin, Genetex #GTX101279; Beta Actin, Genetex #GTX109639; Vinculin, CST #4650S; CHKA, CST #13422S; CDIPT, Genetex #GTX80599; PCYT1A, Sigma-Aldrich #HPA035428; HMOX1, Proteintech #10701-1-AP; FTH1, Cell Signaling #4393S; FLVCR1, Santa Cruz #sc-390100; NDUFS1 Proteintech #12444-1-AP; OPA1 BD Biosciences #612606; MRPS35 Proteintech #16457-1-AP). Secondary antibody incubation was performed at room temperature for 1 hour using anti-mouse IgG-HRP (CST #7076) and anti-rabbit IgG-HRP (CST #7074). Washes were performed with 0.1% Tween-20 tris buffered saline and blots developed using ECL chemiluminescence (Perkin Elmer #NEL105001EA or Cytiva #RPN2232) and autoradiography films (Thomas Scientific #1141J52).

### LipidTOX Staining, Confocal Imaging, and Quantification

Sterile coverslips were coated with 50μg/mL Poly-D-Lysine (ChemCruz #sc-136156) for 1 hour at room temperature after which cells were seeded at a density of 50,000 cells per 6 well. The following day, cells were fixed with 4% paraformaldehyde for 30 minutes at room temperature. Cells were then stained with reconstituted LipidTOX Deep Red (Invitrogen #H34477) diluted 1:1000 in PBS for 30 minutes at room temperature. Coverslips were mounted using Prolong Gold Antifade Mountant (Invitrogen #P10144) and sealed with nail polish. Slides were imaged with a Nikon A1R MP multiphoton microscope with confocal modality, using a Nikon Plan Apo γ 60X/1.40 oil immersion objective. Signal intensity was quantified per cell using FIJI with both mean and maximum fluorescence intensity per cell captured.

### Polar Metabolite Profiling and Isotope Tracing

For in vitro polar metabolomic experiments, 250,000 cells were seeded per 6-well in triplicate per condition. The following day, cells were changed to fresh media, fresh media with vehicle/drug/supplement, or fresh media with 3mg/L [1,2-^13^C_2_]Choline chloride (Cambridge Isotope Laboratories #CLM-548.01) (as indicated in figure legends). 24 hours later, cells were washed twice with ice-cold 0.9% NaCl and polar metabolites extracted in ice cold 80% LC/MS grade methanol containing ^15^N and ^13^C fully-labeled amino acid standards (Cambridge Isotope Laboratories #MSK-A2-1.2). Extracts were vigorously shaken by vortex for 10 minutes at 4°C and spun at 14,000 rpm at 4°C for 10 minutes to remove insoluble cell debris. Supernatants were dried under nitrogen and stored at -80°C until liquid chromatography-mass spectrometry (LC-MS) was performed.

For polar metabolomic experiments from mice, E11.5 embryos were isolated, washed twice in ice cold 0.9% NaCl, and immediately flash frozen with liquid nitrogen. Polar metabolites were extracted in ice cold 80% LC/MS grade methanol containing ^15^N and ^13^C fully-labeled amino acid standards (Cambridge Isotope Laboratories #MSK-A2-1.2). Tissues were homogenized by Bead Ruptor 24 (Omni International) under cooling from liquid nitrogen. Extracts were spun at 14,000 rpm at 4°C for 10 minutes to remove insoluble cell debris. Supernatants were dried under nitrogen and stored at -80°C until liquid chromatography-mass spectrometry (LC-MS) was performed. Insoluble cell debris was used for DNA extraction to identify genotypes of embryos.

LC-MS analysis was conducted on a QExactive benchtop orbitrap mass spectrometer equipped with an Ion Max source and a HESI II Probe coupled to a Dionex Ultimate 3000 UPLC System (Thermo Fisher Scientific). External mass calibration was performed every 3 days using standard calibration mixture.

Dried polar extracts were resuspended in 60 μL of 50% acetonitrile, vortexed for 10 seconds, centrifuged for 15 minutes at (20,000 g, 4°C) and 5 μL of the supernatant was injected onto a ZIC-pHILIC 150 × 2.1mm (5μm particle size) column (EMD Millipore). Chromatographic separation was achieved using the following conditions Mobile phase A consisted of 20 mM ammonium carbonate with 0.1% (v/v) ammonium hydroxide (adjusted to pH 9.3) and mobile phase B was acetonitrile. The column oven and autosampler tray were held at 40°C and 4°C, respectively. The chromatographic gradient was run at a flow rate of 0.150 mL/min as follows: 0-22 min linear gradient from 90% to 40% B; 22-24 min: held at 40% B; 24-24.1 min: returned to 90% B; 24.1-30 min: equilibrated at 90% B.

The mass spectrometer was operated in full-scan, polarity switching mode with the spray voltage set to 3.0 kV, the heated capillary held at 275°C. The sheath gas flow was set to 40 units and the auxiliary gas flow was set to 15 units. The MS1 data acquisition was performed with the following parameters: scan range of 55-825 *m/z*, 70,000 resolution, 1 × 10^6^ AGC target, 80 ms injection time. A pool of all the biological samples was prepared and analyzed using a Top2 data-dependent acquisition method, with polarity switching. The data-dependent MS/MS scans were acquired at a resolution of 17,500, 1 × 10^5^ AGC target, 50 ms max injection time, 1.6 Da isolation width, stepwise normalized collision energy (NCE) of 20, 30, 40 units, 8 sec dynamic exclusion, and loop count of 2.

Relative metabolite abundances were quantified using Skyline Daily v22^58^ using a 3 ppm mass tolerance and a 6 sec retention time from known standards. For *in vitro* experiments, metabolite levels were normalized to protein abundance for each condition as measured by BCA. For *in vivo* experiments, metabolite levels were normalized to endogenous leucine levels.

### Lipid Metabolite Profiling

For *in vitro* polar metabolomic experiments, 250,000 cells were seeded per 6-well in triplicate per condition. The following day, cells were changed to fresh media (as indicated in figure legends). 24 hours later, cells were washed twice with ice-cold 0.9% NaCl and cells lysed in ice cold 80% LC/MS grade methanol. Non-polar metabolites were extracted by the consecutive addition of LC/MS grade water followed by LC/MS grade chloroform. Extracts were vigorously shaken by vortex for 10 minutes at 4°C and spun at 14,000 rpm at 4°C for 10 minutes. The lower lipid-containing layer was carefully collected, dried under nitrogen, and stored at -80°C until LC-MS.

Dried lipid samples were reconstituted in 60 μL 65:30:5 acetonitrile:isopropanol:water (v/v/v), vortexed for 10 sec, centrifuged for 10 min (20,000 g, 4°C) and 5 μl of the supernatant was injected into the LC-MS in a randomized order, with separate injections for positive and negative ionization modes. Lipids were separated on an Ascentis Express C18 2.1 mm x 150mm x 2.7μm column (Supelco) connected to a Dionex UlitMate 3000 UPLC system and a QExactive benchtop orbitrap mass spectrometer (Thermo Fisher Scientific) equipped with a heated electrospray ionization (HESI) probe. Mobile phase A consisted of 10mM ammonium formate in 60:40 water: acetonitrile (v/v) with 0.1% formic acid, and mobile phase B consisted of 10mM ammonium formate in 90:10 isopropanol:acetonitrile (v/v) with 0.1% formic acid. The column oven and autosampler were held at 55°C and 4°C, respectively. The spray voltage was set to 3.5 kV and the heated capillary was held at 320°C. The sheath and auxiliary gas were set to 60 and 20 units, respectively. These conditions were held constant for both positive and negative ionization mode acquisitions. External mass calibration was performed every 3 days using standard calibration mixture.

Mass spectra were acquired, in both positive and negative ionization modes, using a Top15 data-dependent MS/MS method. The full MS scan was acquired as such; 70,000 resolution, 1 × 10^6^ AGC target, 200 ms max injection time, scan range 200 − 2,000 m/z. The data-dependent MS/MS scans were acquired at a resolution of 17,500, AGC target of 1 × 10^5^, 50 ms max injection time, 1.0 Da isolation width, stepwise normalized collision energy (NCE) of 20, 30, 40 units and 8 sec dynamic exclusion.

High-throughput identification and relative quantification of lipids was performed separately for positive and negative ionization mode data using LipidSearch 4.2 Software (Thermo Fisher Scientific/Mitsui Knowledge Industries) with the following search parameters: HCD target database, 5 ppm precursor tolerance, 8 ppm product tolerance, 1% product ion intensity threshold and an m-score threshold of 2. The alignment parameters were: 0.15-0.2 min mean retention time tolerance, A-C ID quality filter, m-score threshold of 5 and the all-isomer peaks node.

After alignment, raw peak areas for all identified lipids were exported to Microsoft Excel. Rejected lipids were removed (‘Rej’ parameter calculated by LipidSearch) and only a single adduct was retained for each lipid. Under these LC-MS conditions, we accepted the following lipid adducts for various classes; sphingolipids: [M+H]/ [M+HCOO]; cholesterol esters, diacyl- and triacylglycerides: [M+NH4]; phosphatidylcholines: [M+H] /[M+HCOO]; phosphoethanolamines: [M+H]/ [M-H]; acylcarnites and monoacylglycerides: [M+H]; other phospholipids (PG, PI):[M+NH4]/ [M-H].

Lipids were then filtered out, if at least one biological group didn’t meet the following criteria: PQ (‘Peak Quality’ parameter calculated by LipidSearch) greater than 0.8; CV (standard deviation/mean peak area) below 0.3; *R* (linear correlation across a three-point dilution series of the representative (pooled) biological sample) greater than 0.8; MS/MS grade of A or B for TG, DG, MG, PE, PG, PI, ChE and HexCer; MS/MS grade of A, B, C for Cer, SM, and PC lipids.

The raw peak areas of the filtered lipids were summed by lipid class for each sample and individual lipid peak areas were normalized to the median lipid signal or protein abundance as measured by BCA for normalization.

### Radioactive Choline Uptake Experiments

Cells were seeded at a concentration of 250,000 or 500,000 cells per 6-well plate in triplicate. The following day, media was aspirated and cells were incubated in room temperature Krebs-Ringer Buffer (Alfa Aesar #J67795) for 30 minutes. Cells were then incubated with indicated concentration of choline chloride in room temperature Krebs-Ringer Buffer for timepoints described in figure legends at room temperature. For all concentrations of choline chloride, 0.093% of the total choline was radioactive ([Methyl-^3^H]-Choline Chloride; Perkin Elmer #NET109001MC)(Choline Chloride; MP Biomedicals #67-48-1). For example, 20nM was radioactive of total 21.49μM choline used in the majority of experiments. Following incubation, cells were washed twice with ice cold Krebs-Ringer Buffer on ice. Cells were then solubilized with 200μL of 1% SDS 0.2N NaOH and transferred to a scintillation vial with 5mL of Insta-Gel Plus scintillation cocktail (Perkin Elmer #601339). Radioactivity was measured with the TopCount scintillation counter (Perkin Elmer). Radioactivity was normalized to the seeded cell number for each experiment.

For some experiments, Krebs-Ringer Buffer without Sodium (0.54mM KCl, 1.3mM CaCl_2_, 0.53mM MgCl_2_, 0.4mM MgSO_4_, 0.37mM KH_2_PO_4_, 240mM sucrose, 4.4mM Tris-phosphate, 5.5mM Glycine, 5.6mM D-Glucose; pH 6.5 with KOH) or Krebs-Ringer Buffer with Sodium (0.54mM KCl, 1.3mM CaCl_2_, 0.53mM MgCl_2_, 0.4mM MgSO4, 0.37mM KH_2_PO_4_, 138mM NaCl, 0.28 mM Na_2_HPO_4_, 5.5mM Glycine, 5.6mM D-Glucose; pH 6.5 with KOH) were used. In these experiments, all incubations and washes used the same Krebs-Ringer Buffer.

For the hemicholinium-3 competition experiments, hemicholinium-3 was serially diluted in Krebs-Ringer Buffer with radioactive choline and cells were treated with both compounds at the same time.

### Rapid Mitochondrial Purification for Metabolite Profiling and Proteomics

Mitochondria were purified from HeLa FLVCR1 KO and FLVCR1 KO +FLVCR1 cDNA cells expressing a 3xHA-OMP25-mCherry (mitochondrial isolation) or 3xMyc-OMP25-mCherry (background control) construct. Cells were cultured in choline depleted or choline replete media for 2 weeks prior to this experiment. Equal cell number was seeded in 15cm plates and media was changed to choline depleted or choline replete media with 1% dFBS 24 hours prior to mitochondrial isolation. Per condition, triplicate confluent 15cm plates were washed twice with ice cold KPBS and scraped in 1mL of ice cold KPBS. Cells were pelleted via centrifugation at 1,000*g* for 1.5 minutes at 4°C. Cells were resuspended in 1mL of ice cold KPBS and homogenized with 30 passes of a 2mL dounce homogenizer. Homogenate was spun at 1,000*g* for 1.5 minutes at 4°C to pellet intact cells. 5μL of cleared homogenate was added to 45μL of TritonX Lysis Buffer as a whole cell lysate sample and 20μL of cleared homogenate was added to 120μL of methyl tert-butyl ether (MTBE) for extraction of whole cell lipids. The remaining homogenate was incubated with 200μL of prewashed anti-HA magnetic beads (Thermo Scientific #88837) on a rotator for 5 minutes at 4°C. Following incubation, beads were washed three times with ice cold KPBS. Half of the beads were incubated with 90μL TritonX Lysis Buffer for proteomics while the other half were incubated with 240μL of MTBE for lipidomics on a rotator for 10 minutes at 4°C. For proteomic samples - following incubation, eluate was collected from beads and spun at 1,000*g* for 1.5 minutes at 4°C to remove cellular debris and potential bead contamination. 5μL of supernatant was added to 45μL of TritonX Lysis Buffer as a mitochondrial lysate sample. The remaining supernatant was used for quantitative proteomics and stored at -80°C until LC-MS-MS. For lipidomic samples – following incubation eluate was collected from beads and added to 200μL of ice-cold LC/MS grade water. 100μL of ice-cold LC/MS grade water was also added to whole cell lipidomic samples at this point. Both whole cell and mitochondrial lipidomic samples were vigorously shaken by vortex for 10 minutes at 4°C and spun at 14,000 rpm at 4°C for 10 minutes. The lower lipid-containing layer was carefully collected, dried under nitrogen, and stored at -80°C until LC-MS.

### Mitochondrial Proteomics

Cysteines were reduced and alkylated using DTT and IAA. Proteins were precipitated with acetone and pellets were digested with trypsin in 200mM EPPS. Peptides were labeled with TMTpro aliquots, quenched with hydroxylamine and pooled. Pooled peptides were fractionated using high pH reverse phase spin columns (Pierce) and analyzed by LC-MSMS with MS2 fragmentation. Data was processed using Proteome Discoverer v. 2.5 and further statistical analysis was performed within the Perseus statistical software environment^59^. All values were log_2_ transformed and normalized to the median intensity within each sample. Proteins with at least three detected peptides were filtered for further analysis. A truncated ANOVA test based on this criterion and a permutation-based FDR q<0.01 identified 411 proteins which were used to generate the principal component analysis (PCA) and heatmap.

### RNA-Seq

Cells were seeded at a concentration of 250,00 cells per 6-well in triplicate in choline depleted or replete medium. The next day cells were given fresh media and 24 hours later collected for RNA extraction using the RNeasy Mini Kit (Qiagen #74104) according to manufacturer’s manual. RNA concentrations were determined using Qubit 2.0 Fluorometer (Life Technologies) and RNA integrity assessed by Agilent TapeStation 4200 (Agilent Technologies). RNA sequencing libraries were prepared using the NEBNext Ultra RNA library kit for Illumina (NEB) following manufacturer’s specifications. Sequencing libraries were clustered onto a single lane of a flowcell and loaded on a Illumina HiSeq Instrument according to manufacturer’s guidelines. Samples were sequenced using a 1×75bp Single End configuration. Sequencing reads were aligned to the reference genome (Homo_sapiens.GRCh38) and gene models retrieved from the TxDB.Hsapiens.UCSC.hg38.knownGene.gtf.gz Bioconductor library. Transcript expression was calculated using Salmon^60^ and raw read counts per gene were normalized and rlog transformed using DESeq2^61^. Significant differentially expressed genes between conditions were identified by DESeq2with a Benjamini Hochberg adjusted p-value <0.05. Geneset Enrichment Analysis (GSEA) was performed using the fgsea() R package^62^.

### Proliferation Assays

HeLa and HEK293T *FLVCR1* KO and *FLVCR1* KO +*FLVCR1* cDNA cells were seeded at 1.5×10^6^ cells per 15cm plate in either choline depleted or choline replete media. Cells were maintained in media conditions and passaged together by cell line as needed with 1.5×10^6^ reseeded at each passage. With each passage, cell number was counted in triplicate and cumulative log_2_ population doublings calculated.

After passage in respective medias for at least 1 week, cells were seeded in 96-well plates at a concentration of 1,000 cells per well in a final volume of 200μL with indicated media conditions. Cells were seeded in at least triplicate wells per condition. For initial cell number of each condition, 40μL of Cell Titer Glo Reagent (Promega) was added and luminescence read using a SpectraMax M3 plate reader (Molecular Devices) on the day of seeding, After 5 (HEK293T) or 6 (HeLa) days of growth, luminescence was read as above. Data are presented as log_2_fold change in cell number relative to luminescence of initial cell number of each condition.

### CRISPR/Cas9 Genetic Screens

Metabolism-scale CRISPR knockout screens were performed as previously described^38,63^. Briefly, 2,989 genes encoding metabolic enzymes and small molecular transporters were targeted with a total of 23,777 sgRNAs and 50 non-targeting sgRNA controls. After cloning into lentiCRISPR-v2 puro (Addgene #982990), the pooled plasmid library was used to produce lentivirus-containing supernatants. The titer of lentiviral supernatants was determined by infection of target cells at a range of virus and counting the number of puromycin-resistant cells after 3 days of selection. For the screens, 30×10^6^ target cells were infected at an MOI of ∼0.7 and selected with puromycin. An initial pool of 30×10^6^ cells was harvested for genomic DNA extraction at the beginning of the screen. 30×10^6^ cells at a concentration of 3×10^6^ per 15cm plate were subjected to each experimental condition and passaged every 3 days until ∼14 cumulative population doublings were reached. The following target cells and conditions were used for the screens in this study – HEK293T FLVCR1 KO cells: RPMI 1640, RPMI 1640 +1mM Choline, Choline Depleted RPMI 1640, Choline Replete RPMI 1640; HEK293T FLVCR1 KO +FLVCR1 cDNA cells: RPMI 1640. On the final day of screening, cells were harvested for genomic DNA extraction. Genomic DNA was extracted using a DNeasy Blood & Tissue Kit (Qiagen) and amplification of sgRNA inserts was performed via PCR using barcoded primers for each condition. PCR amplicons were purified and sequenced on a MiSeq (Illumina). Sequencing reads were mapped and the abundance of each sgRNA was measured. Gene score is defined as the median log_2_fold change in the abundance of all sgRNAs targeting a particular gene between the initial and final populations.

### Mice

All animal studies were performed according to a protocol approved by the Institutional Animal Care and Use Committee (IACUC) at The Rockefeller University. Animals were housed in ventilated caging on a standard light/dark cycle with food and water provided *ad libitum. Flvcr1*^HET^ mice backcrossed to a C57BL/6 background were kindly provided by Janis Abkowitz and previously described^9^. Of note, exon 3 of *Flvcr1* is deleted in these mice. As reported, we did not identify any *Flvcr1*^KO^ animals born. Previous reports suggest that *Flvcr1*^KO^ mice in a pure C57BL/6 genetic background die at around E12.5. We set up timed matings of *Flvcr1*^HET^ mice and staged embryos according to standard methods with the day of a copulatory plug identified as 0.5dpc. E11.5 embryos were isolated with the aid of a stereo microscope (Zeiss) and imaged. Whole embryos were used for polar metabolite profiling and western blotting. The following primers were used to genotype mice with wild-type alleles producing a PCR product at 374bp and knockout alleles producing a PCR product at 674bp: P1R: 5’-CAATAGACATTTAACACCCC-3’; P2F: 5’-CAAGAGTTCTATCTGGAACC-3’; P3F3: 5’-CCTGCCTTGAGATAGCTGC-3’.

### METSIM Dataset Analysis

We compiled a list of ∼3000 genes of interests that included a union set of metabolic genes and plasma membrane transporters. The boundaries of the genes were defined according to the ENSEMBL reference human genome sequence (Homo_sapiens.GRCh38.106.gtf) ±15 kB pairs upstream and downstream. Given the specified window, we applied the minimum-p-value gene-based approach to represent the metabolite-gene associations from the METSIM dataset that contains the GWAS results for blood serum level of 1391 metabolites from 6136 men^4^. Then, metabolite-wise and gene-wise ranks were computed based on the observed p-value statistic (with the top rank 1 corresponding to the lowest p-value statistic or strongest association) We note that an empirical p-value can be computed using permutation, as the limiting distribution of the gene-based statistic is not known under the null hypothesis that a gene is not associated with a metabolite. To filter out top associations, we defined the following criteria all of which must be satisfied: (i) the gene must be ranked in top 10 for a given metabolite; (ii) metabolite must be the top score for the considered gene; (iii) -log10(p-val) > 5 of the association (defined by Bonferroni multiple test correction). Afterwards, based on the overlap between AMIGO GO Term “Plasma Membrane” and an in-house transporter gene list, plasma membrane transporters were selected for further analysis. The resulting entries were distributed in four categories: (a) metabolites are classified as xenobiotics/partially characterized/uncharacterized, (b) previously reported in literature associations, (c) unknown associations with metabolites that score only for one plasma membrane transporter, (d) other entries. Metabolite classification followed that used in the METSIM study^4^.

### DepMAP and METSIM Integrated Analysis

METSIM associations were identified through the minimum-p-value approach and ranked as described above. The top associations were defined by the following criteria: (i) the gene must be ranked in top 10 for a given metabolite; (ii) -log10(p-val) > 5 of the association (defined by Bonferroni multiple test correction). Then, entries containing xenobiotics/partially characterized/uncharacterized metabolites were removed. Pearson Correlation Coefficient (PC) values were computed for the selected list of ∼3000 metabolic genes from DepMap 22Q2 Public+Score, Chronos dataset. Then, we filtered interactions containing plasma membrane transporters with |PC| > 0.2. To avoid false positive hits, we removed the entries in which both genes are located on the same chromosome. Finally, to integrate METSIM and DepMap, we selected the pairs with |PC| > 0.2 in which both genes shared the same top-scoring metabolite according to our METSIM analysis.

