## Supplementary figures and images for "Integrative genetic analysis identifies FLVCR1 as an essential component of choline transport in mammals"

### Supplemental Figure 1

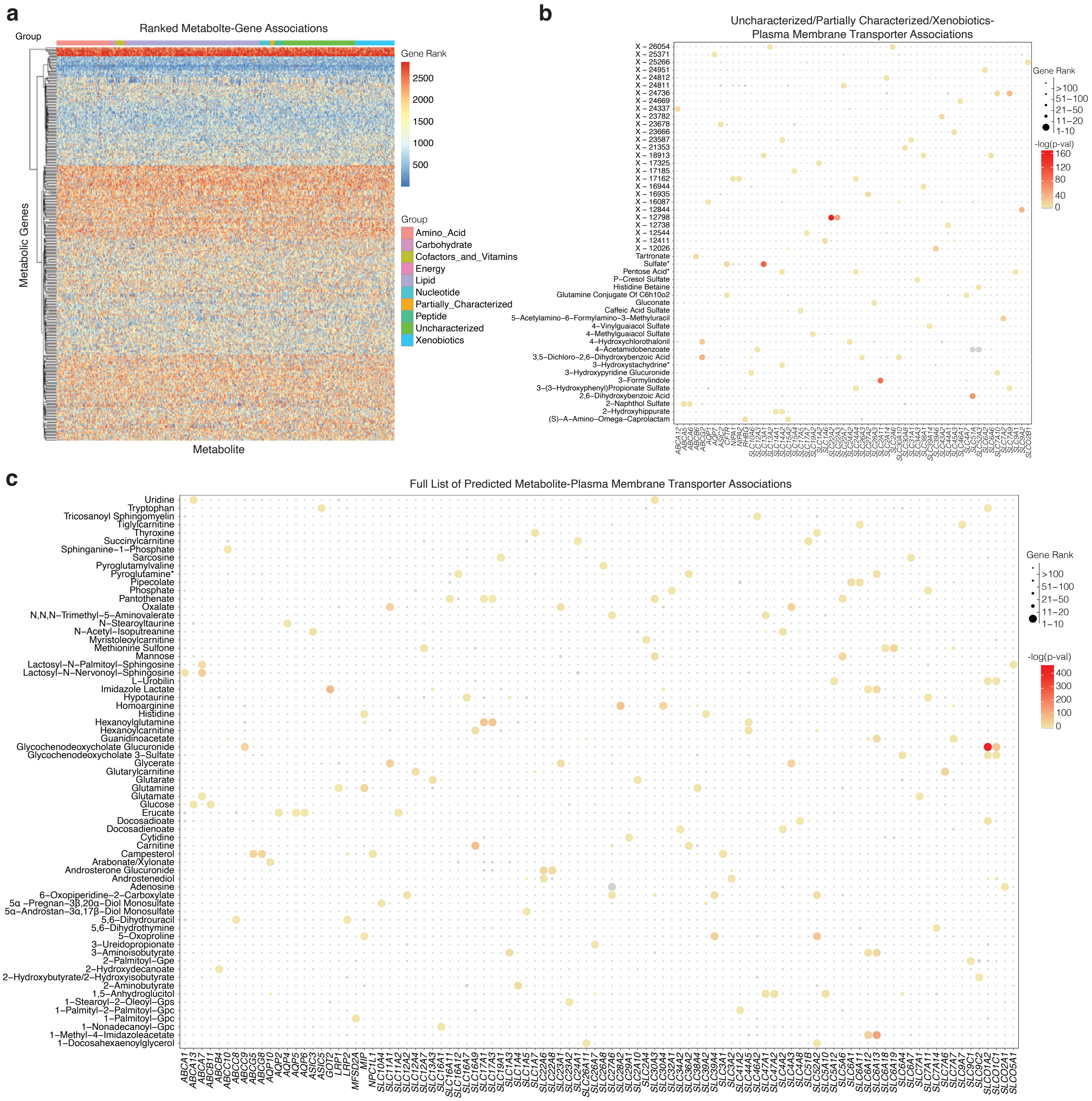

### Supplemental Figure 2

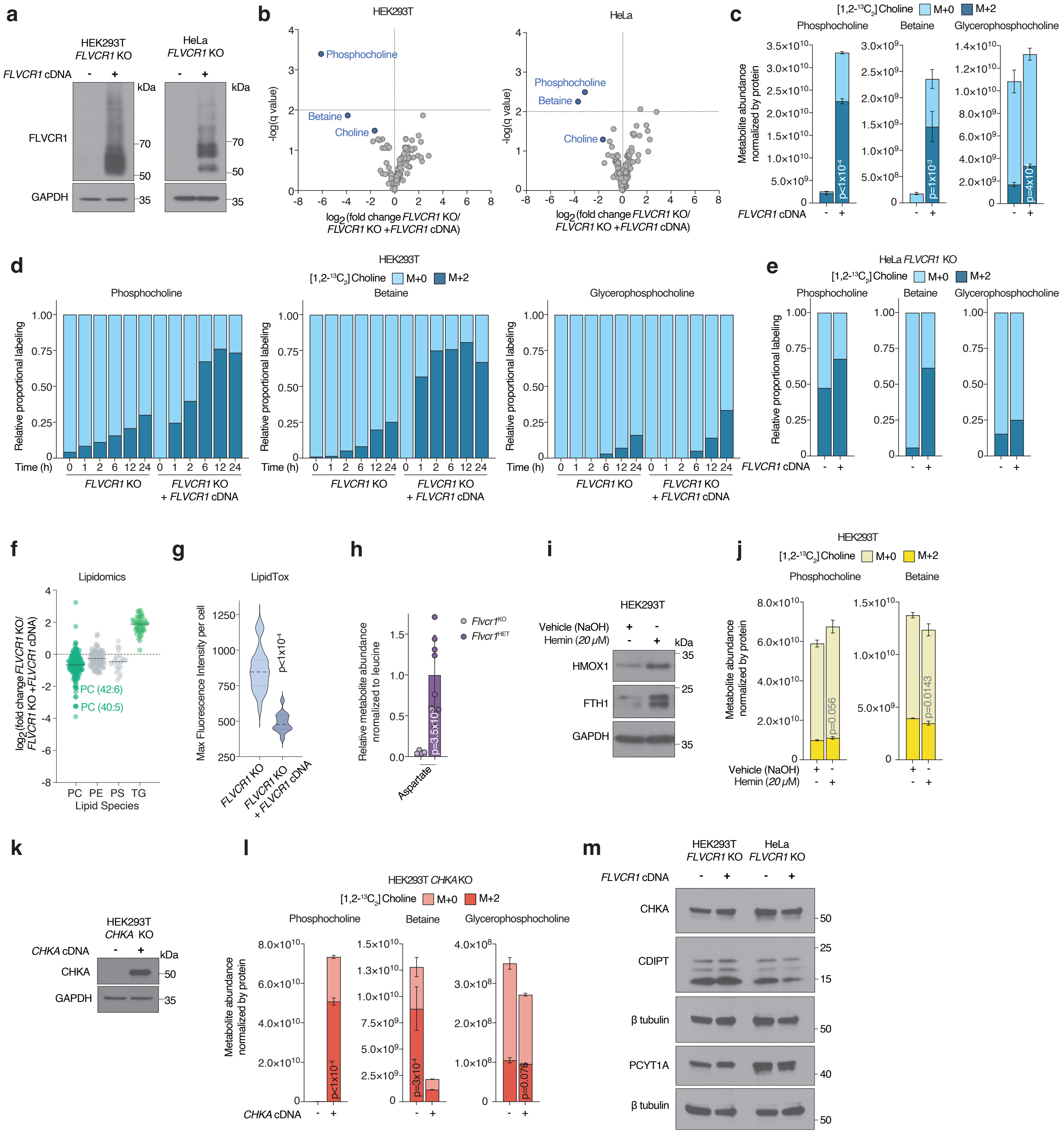

### Supplemental Figure 3

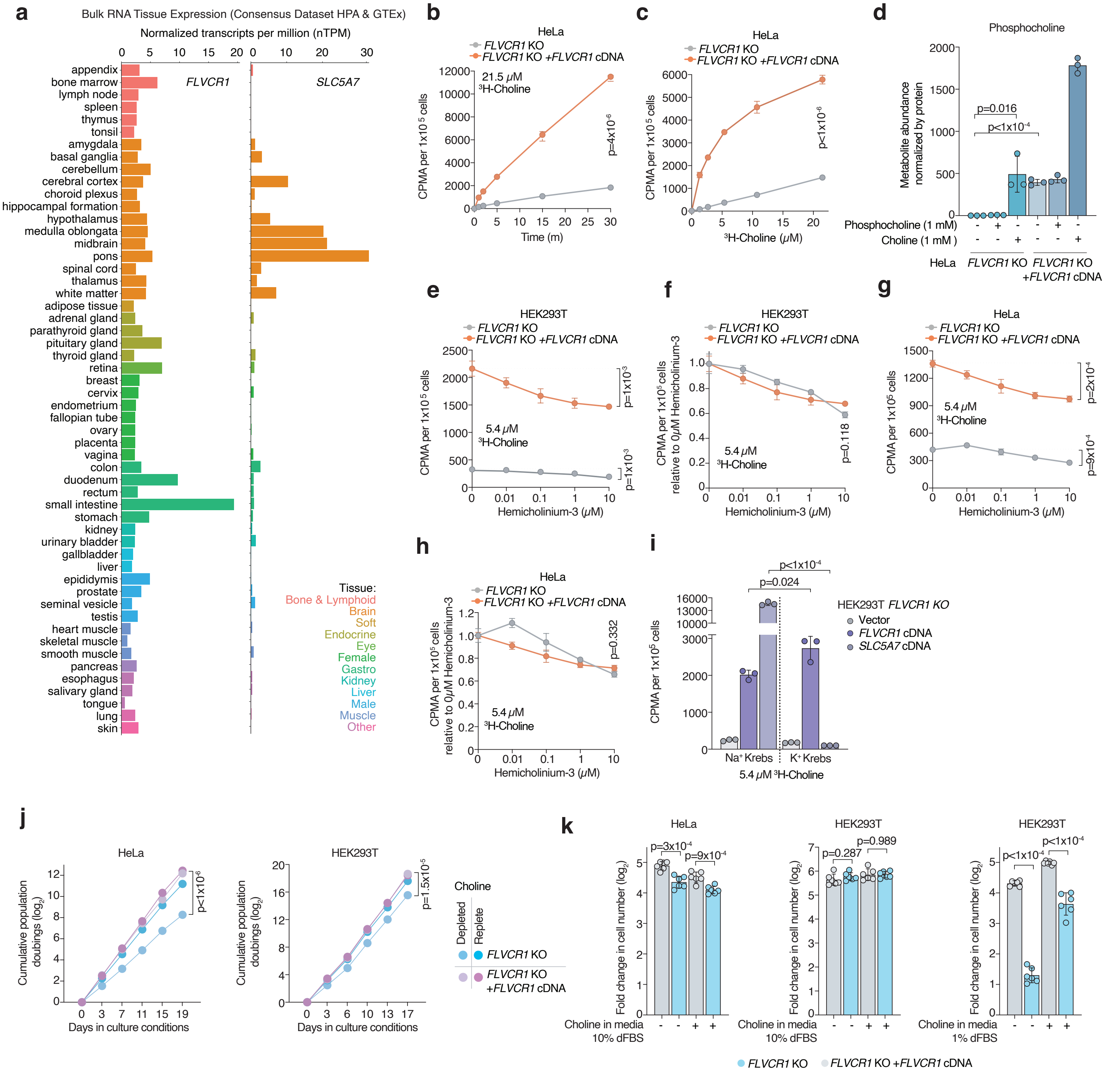

### Supplemental Figure 4

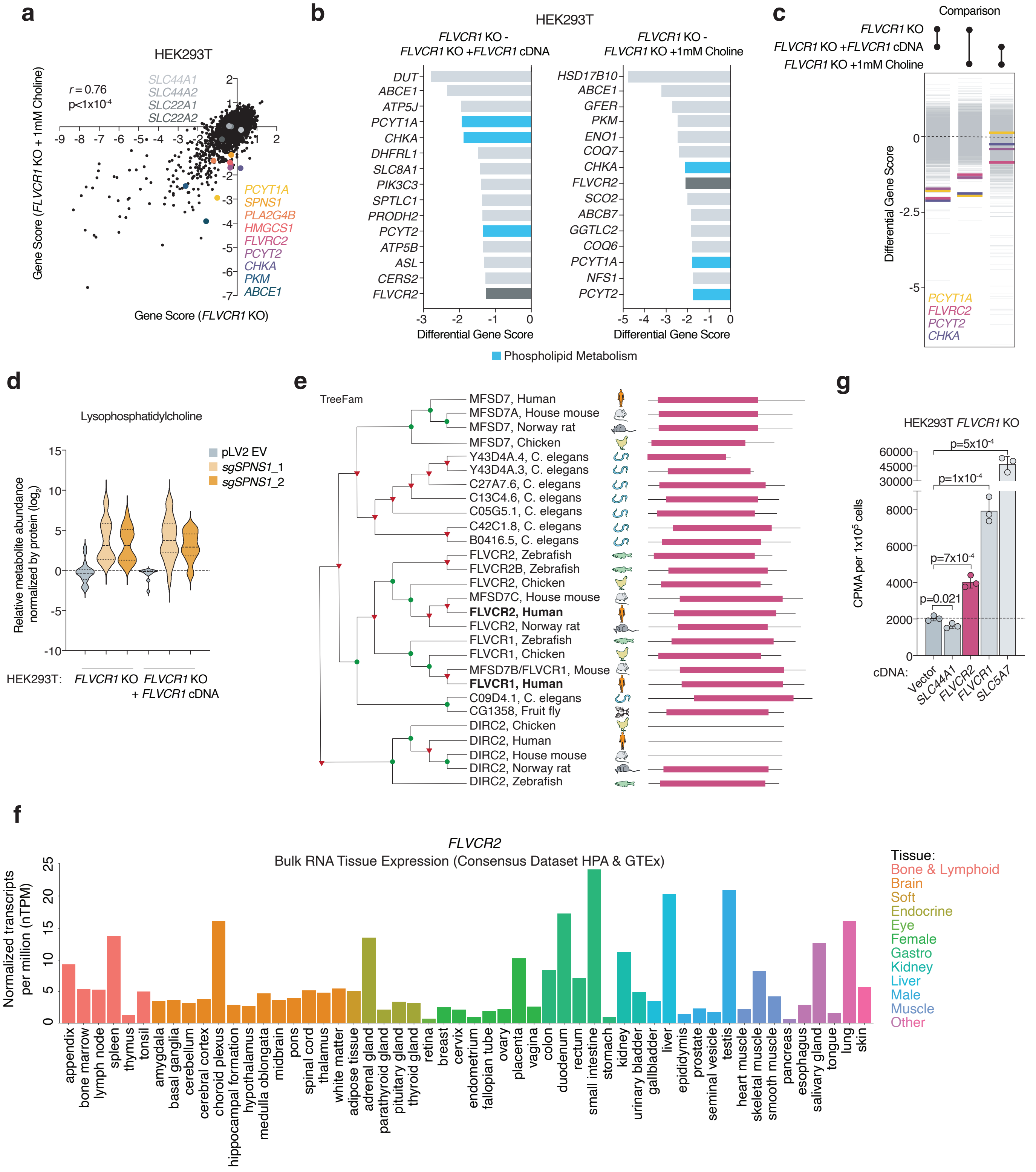

### Supplemental Figure 5

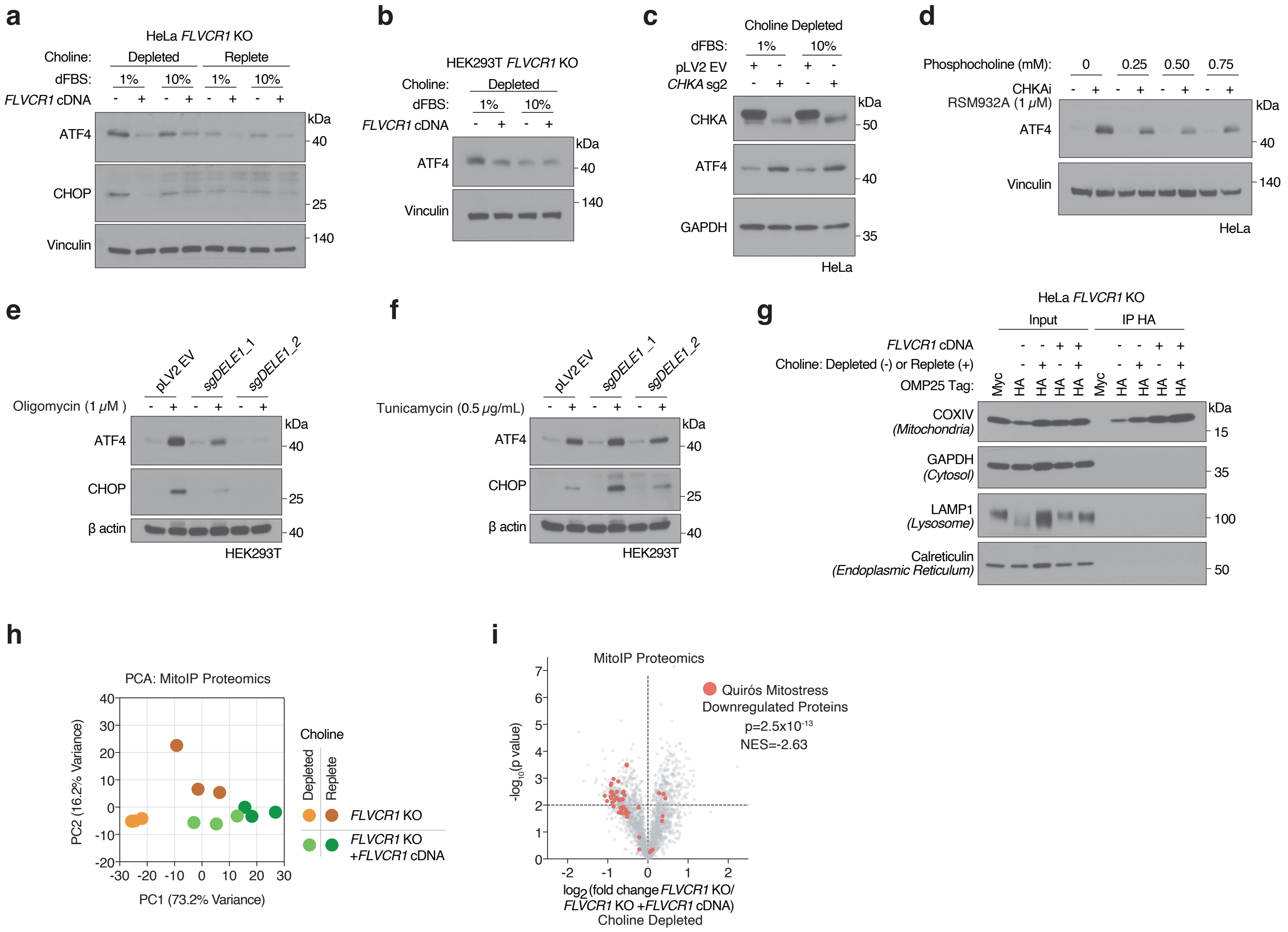
